# A synergistic core for human brain evolution and cognition

**DOI:** 10.1101/2020.09.22.308981

**Authors:** Andrea I. Luppi, Pedro A.M. Mediano, Fernando E. Rosas, Negin Holland, Tim D. Fryer, John T. O’Brien, James B. Rowe, David K. Menon, Daniel Bor, Emmanuel A. Stamatakis

## Abstract

A fundamental question in neuroscience is how brain organisation gives rise to humans’ unique cognitive abilities. Although complex cognition is widely assumed to rely on frontal and parietal brain regions, the underlying mechanisms remain elusive: current approaches are unable to disentangle different forms of information processing in the brain. Here, we introduce a powerful framework to identify synergistic and redundant contributions to neural information processing and cognition. Leveraging multimodal data including functional MRI, PET, cytoarchitectonics and genetics, we reveal that synergistic interactions are the fundamental drivers of complex human cognition. Whereas redundant information dominates sensorimotor areas, synergistic activity is closely associated with the brain’s prefrontal-parietal and default networks; furthermore, meta-analytic results demonstrate a close relationship between high-level cognitive tasks and synergistic information. From an evolutionary perspective, the human brain exhibits higher prevalence of synergistic information than non-human primates. At the macroscale, we demonstrate that high-synergy regions underwent the highest degree of evolutionary cortical expansion. At the microscale, human-accelerated genes promote synergistic interactions by enhancing synaptic transmission. These convergent results provide critical insights that synergistic neural interactions underlie the evolution and functioning of humans’ sophisticated cognitive abilities, and demonstrate the power of our widely applicable information decomposition framework.

## Synergistic and redundant interactions identify brain networks with distinct neurocognitive profiles

In theoretical and cognitive neuroscience, considering the human brain as a distributed information-processing system has proven to be a powerful framework to understand the neural basis of cognition ^1^. Crucially, a deeper understanding of any information-processing architecture calls for a more nuanced account of the information that is being processed.

As an example, let us consider humans’ two main sources of information about the world: the eyes. The information that we still have when we close either eye is called “redundant information” — because it is information that can be conveyed by either source (for instance, information about colour is largely redundant between the two eyes). Redundancy provides robustness: we can still see with one eye closed. However, closing one eye also deprives us of stereoscopic information about depth. This information does not come from either eye alone: ones needs both, in order to perceive the third dimension. This is called the “synergistic information” between two sources - the extra advantage that we derive from combining them, which makes them complementary ^2,3^.

Thus, in addition to their own unique information, when multiple sources are considered together their information contribution can be identified as synergistic (only available when both sources are considered together) or redundant (available from either source independently). Every information-processing system — including the human brain — needs to strike a balance between these mutually exclusive kinds of information, and the advantages they provide: robustness and integration, respectively ^4–7^. Being fundamentally different, synergistic and redundant information cannot be adequately captured by traditional measures of macroscale information exchange (“functional connectivity”) in the human brain, which instead simply quantify the similarity between regional activity ^2,8^.

Here, we reveal the distinct contributions of synergistic and redundant interactions to human cognition, and we delineate their large-scale organisation in the human brain. To this end, we leveraged the partial information decomposition (PID) framework ^2,3,9^ to quantify synergistic and redundant interactions between brain regions (Figure 1A,B), obtained from resting-state functional MRI data from 100 Human Connectome Project subjects (Methods). We ranked each brain region separately in terms of how synergistic and redundant its interactions with other brain regions are; the difference between these ranks (synergy minus redundancy) determines the relative relevance of a given region for synergistic versus redundant processing, thereby defining a redundancy-to-synergy gradient across brain regions (Figure 1C).

**Figure 1.**
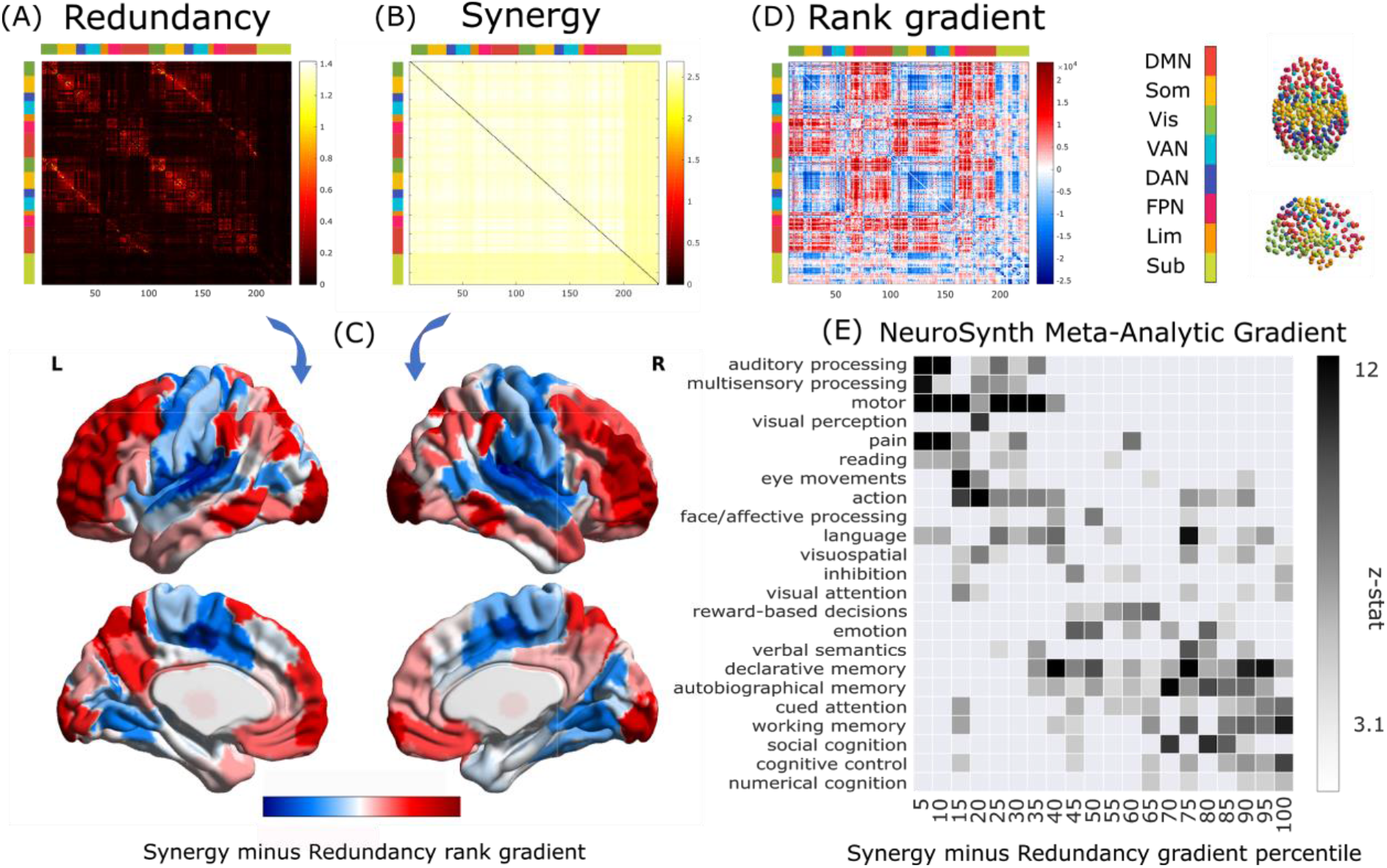
Synergistic and redundant networks exhibit distinct anatomical and cognitive profiles. Group-average matrices of redundant (A) and synergistic (B) interactions between regions of the 232-ROI augmented Schaefer atlas. (C) Brain surface projections of regional redundancy-to-synergy gradient scores, obtained as the difference between each region’s rank in terms of synergy and in terms of redundancy; positive scores (red) indicate a bias towards synergy, and negative scores (blue) a bias towards redundancy. (D) Matrix of redundancy-to-synergy gradient scores (synergy minus redundancy ranks) for each connection between brain regions. (E) Results of the NeuroSynth term-based meta-analysis, relating the distribution of redundancy-to-synergy gradient across the brain (discretised in 5% increments) to a gradient of cognitive domains, from lower-level sensorimotor processing to higher-level cognitive tasks. These results are robust to the use of different parcellations (cortical-only, having lower or higher number of nodes, and obtained from anatomical rather than functional considerations; Figure S1A-C) and are also replicated without deconvolving the hemodynamic response function from the functional data (Figure S1D).

Our results demonstrate that traditional FC mostly captures redundant, rather than synergistic, information exchange in the human brain (Figure S2). Furthermore, they clearly show that redundant and synergistic interactions delineate networks with distinct neuroanatomical profiles (Figure 1A-D). In terms of Von Economo’s cytoarchitectonic classification ^10^, redundant interactions are especially prominent in primary sensory, primary motor and insular cortices (Figure S3), corresponding to the brain’s somatomotor and salience subnetworks (Figure S4). In contrast, regions with higher relative importance for synergy predominate in higher-order association cortex, and are affiliated with the default mode (DMN) and fronto-parietal executive control (FPN) subnetworks ^11^ (Figures S3-4).

It is noteworthy that synergy, which quantifies the extra information gained by integrating multiple sources ^3,12^ is most prevalent in regions belonging to the DMN and FPN. Functionally, these regions are recruited by complex tasks that rely on multimodal information, decoupled from immediate sensorimotor contingencies ^13,14^; anatomically, they receive multimodal inputs from across the brain ^15^. Therefore, it has been speculated that these networks are devoted to the integration of information ^13,15^. Our findings about regional prevalence of synergy in DMN and FPN provide formal information-theoretic evidence to confirm this long-standing hypothesis. Furthermore, by considering a synergy-redundancy gradient in terms of connections instead of regions, we show that the most synergy-dominated connections correspond to links between DMN/FPN and other subnetworks, whereas redundancy-dominated connections tend to occur within each subnetwork (Figure 1C).

The distinct cytoarchitectonic profiles and subnetwork affiliations further suggest that redundant and synergistic interactions may be involved with radically different cognitive domains. To empirically validate this hypothesis, we performed a term-based meta-analysis using NeuroSynth. The redundancy-to-synergy gradient identified in terms of regional rank differences was related to 24 terms pertaining to higher cognitive functions (e.g. attention, working memory, social and numerical cognition) and lower sensorimotor functions (such as eye movement, motion, visual and auditory perception) adopted by previous studies ^13,16^.

Supporting the inference from neuroanatomy to cognition, our results reveal that the regional gradient from redundancy to synergy corresponds to a gradient from lower to higher cognitive functions. Specifically, high-redundancy regions loaded strongly onto auditory, visual and multisensory processing and motion. In contrast, high-synergy regions had the strongest loadings onto social and numerical cognition, working memory and cognitive control (Figure 1E).

## Network organisation of synergy and redundancy support their distinct information-processing roles

Sensorimotor and higher-order cognitive functions impose distinct and opposite demands on cognitive architectures: specialised sensory processing benefits from segregation into modules, whereas integration of information demands high levels of interconnectedness ^5,17^. Contrasting the properties of the networks delineated by synergistic and redundant interactions reveals how the human brain resolves this tension.

Across individuals, the network of synergistic interactions is more highly interconnected and globally efficient than the network of redundancy (Synergy: M=2.54, SD=0.06; Redundancy: M=0.14, SD=0.04; t(99)=-330.04, p<0.001, Hedge’s *g=-*46.67) (Figure 2A). In contrast, redundant interactions delineate a network characterised by a highly modular structure, which is virtually absent in synergistic networks (Synergy: M=0.005, SD=0.001; Redundancy: M=0.29, SD=0.06; t(99)=51.74, p<0.001, Hedge’s *g=7*.*25*) (Figure 2B). Thus, synergistic and redundant interactions exhibit distinct network organisation, supporting integrated and segregated processing, respectively - as demanded by the cognitive functions they support.

**Figure 2.**
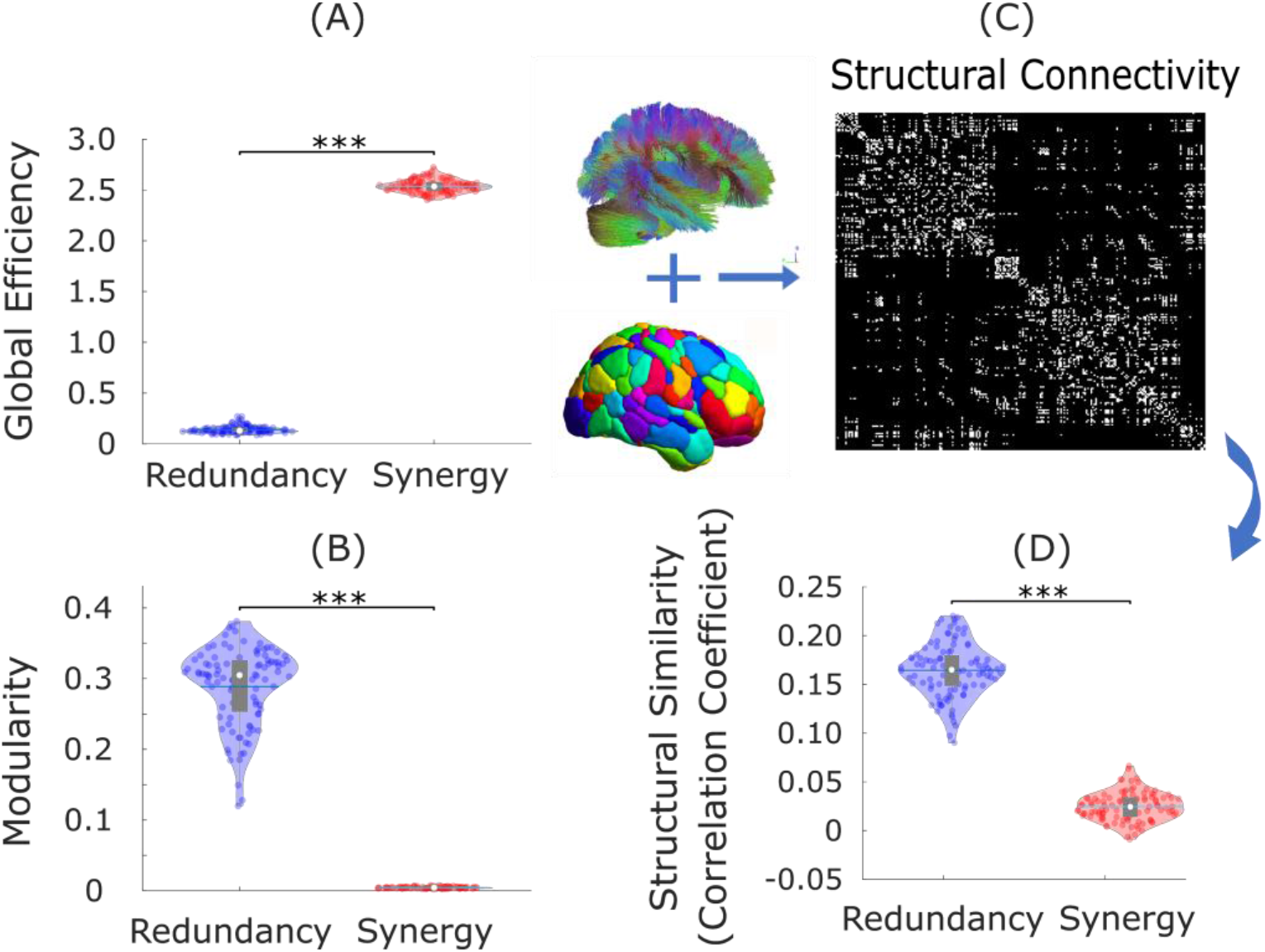
Synergy is integrated, redundancy is segregated and supported by anatomical connections. (A) The network organisation of synergistic interactions exhibits significantly higher integrative capacity (global efficiency) than redundant interactions. (B) The network organisation of redundant interactions exhibits significantly higher segregation (modularity) than synergistic interactions. (C) Structural connectivity of each subject was estimated from diffusion MRI, measured as the number of white matter tracts between regions of the 232-ROI augmented Schaefer atlas, and Spearman correlation coefficient was used to assess the similarity of redundancy and synergy matrices with structural connectivity, after thresholding to ensure equal numbers of connections. (D) Networks of redundant interactions are significantly more correlated with underlying structural connectivity than synergistic interactions. Violin plots represent the distribution of values across 100 HCP subjects (colored circles). White circle: mean; blue line: median; grey box: interquartile range; *** p < 0.001.

It is also known that only a subset of regions are directly connected by white matter tracts ^18^; therefore, we reasoned that the more an organism’s survival depends on information exchange between regions X and Y, the more one should expect X and Y to be directly connected. Thus, direct physical connections in the brain reveal where the need for robust communication is highest. Consequently, if redundant interdependencies are representative of robust information exchange, they should be co-located with underlying direct anatomical connections - as quantified using diffusion-weighted imaging (DWI). Our results support this hypothesis: across subjects, the number of white matter streamlines was significantly more correlated with redundant (M=0.16, SD=0.028) than synergistic interactions between regions (M=0.025, SD=0.015; t(99)=39.85, p<0.001, Hedge’s *g=*6.29) (Figure 2C,D). These results are replicated using alternative network measures and parcellations (Figures S5-7 and Supplementary Tables 1-3).

Thus, whereas synergistic interactions are poised to facilitate high-level cognition through global integration, redundant interactions demarcate a structural-functional backbone in the human brain, ensuring robust sensorimotor input-output channels - both critical functions for successful information processing.

## High-synergy brain regions are selectively potentiated by human evolution

The association between synergistic information processing and higher cognitive functions, raises the intriguing possibility that the human brain may enable humans’ uniquely sophisticated cognitive capacities in virtue of its highly synergistic nature. We pursued this hypothesis through three convergent approaches.

First, we show that the human brain is especially successful at leveraging synergistic information, compared with the brains of non-human primates. Synergistic interactions account for a higher proportion of total information exchange in the human brain than in the macaque (Macaca mulatta); whereas the two species’ brains are equal in terms of proportion of total information exchange accounted for by redundancy (Synergy: Human M=0.478, SD=0.003; Macaque M=0.466, SD=0.005; t(117)=14.24, p<0.001, Hedge’s g=3.54; Figure 3A; Redundancy: Human M=0.012, SD=0.005; Macaque M=0.011, SD=0.005; t(117)=0.90, p=0.372, Hedge’s g=0.22; Figure 3B).

**Figure 3.**
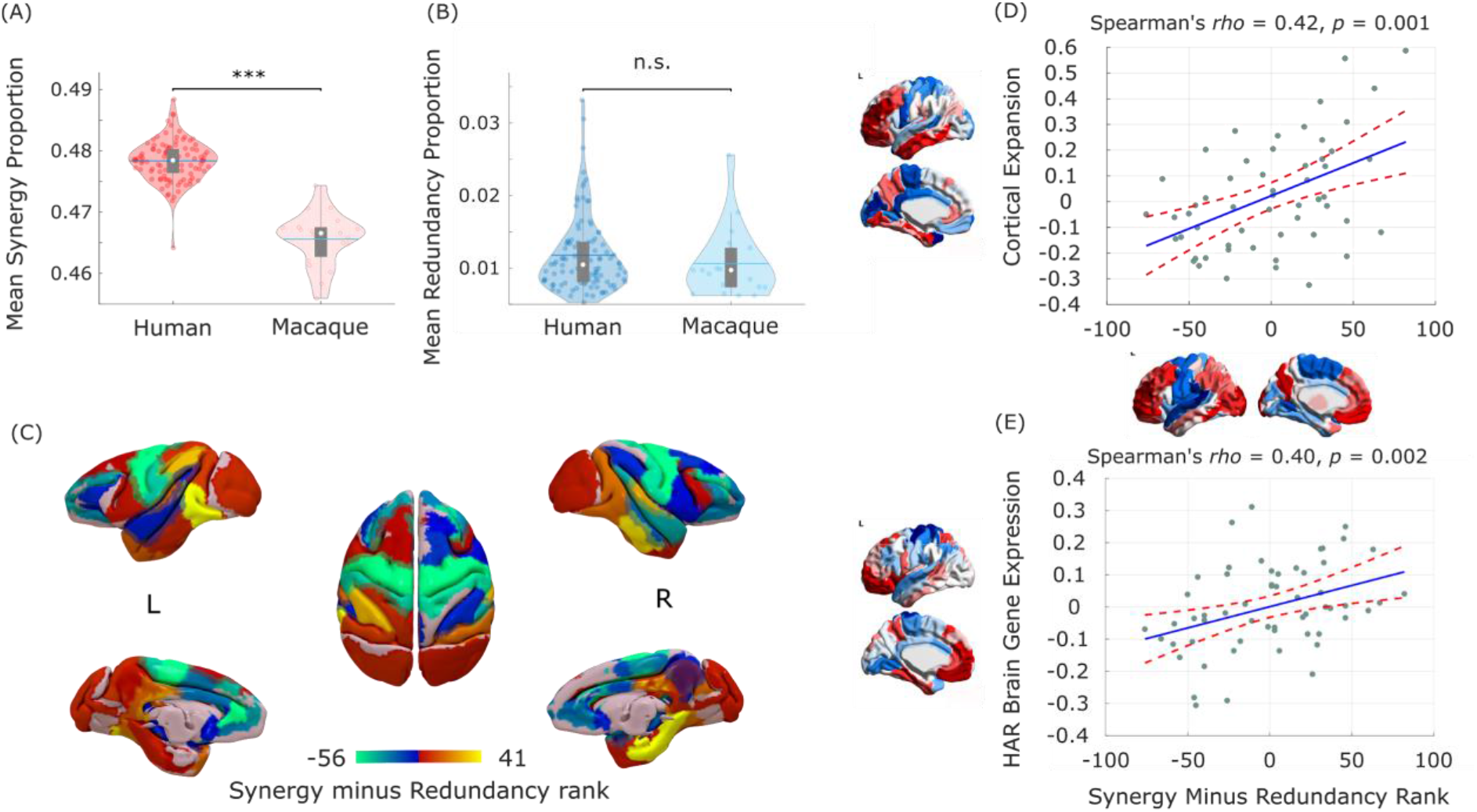
Human brain evolution favoured high synergy. (A) The proportion of synergistic information exchange across the brain is significantly higher in humans (Homo sapiens) than macaques (Macaca mulatta). (B) The proportion of redundant information exchange across the brain is equivalent in humans and macaques. (C) Surface projection of regional redundancy-to-synergy gradient scores for the macaque brain. (D) Significant correlation between human regional redundancy-to-synergy gradient scores and regional cortical expansion from chimpanzee (Pan troglodytes) to human (both on left hemisphere of DK-114 cortical atlas). (E) Significant correlation between human regional redundancy-to-synergy gradient scores and regional expression of brain-related human-accelerated (HAR-Brain) genes (both on left hemisphere of DK-114 atlas). The results in (A) and (B) cannot be solely attributed to either the choice of bandpass filter, or the difference in TR between datasets (Figures S9-10). The results in (D) and (E) are also replicated using unadjusted scores (Figure S11).

The patterns of synergy and redundancy in the macaque brain broadly resemble those observed in humans (Figure S8 and Supplementary Table 7), demonstrating their evolutionary stability - including the expected high redundancy in sensorimotor regions (Figure 3C). However, redundancy is more prevalent than synergy in the prefrontal cortex (PFC) of macaques, despite PFC being among the most synergy-dominated cortices in humans (Figure 3C). Intriguingly, prefrontal cortex underwent substantial cortical expansion in the course of human evolution ^19^.

These findings suggest that the high synergy observed in human brains may be a specific outcome of evolutionary cortical expansion. To explore this hypothesis, we analysed cortical morphometry data from in vivo structural MRI, comparing humans and one of the closest evolutionary relatives of Homo sapiens: chimpanzees (Pan troglodytes)^20^. Supporting our hypothesis, we identified a significant positive correlation between relative cortical expansion in humans versus chimpanzees, and the gradient of regional prevalence of synergy previously derived from functional MRI (ρ = 0.42, p = 0.001; Figure 3D). Thus, these findings suggest that the additional cortical tissue gained through human evolution is primarily dedicated to synergy, rather than redundancy.

To provide further support for the evolutionary relevance of synergistic interactions, we capitalised on human adult brain microarray datasets across 57 regions of the left cortical mantle ^20^, made available by the Allen Institute for Brain Science (AIBS) ^21^. We demonstrate that regional dominance of synergy correlates with regional expression of genes that are both (i) related to brain development and function, including intelligence and synaptic transmission ^20^; and (ii) selectively accelerated in humans versus non-human primates (“HAR-Brain genes”; ρ = 0.40, p = 0.002; Figure 3E). Thus, the more important a brain region is in terms of synergy, the more likely it is to express brain genes that are uniquely human.

Taken together, these findings provide converging evidence for the hypothesis that evolutionary pressures selectively potentiated the role of synergistic interactions in the human brain, both in terms of dedicated genes, (Fig. 3E) dedicated cortical real estate (Fig. 3D), and the end result: higher prevalence of synergy in human brains than non-human primates (Fig. 3A,B).

## Neurobiological origins of synergy in the human brain

These observations raise the question of how such high synergy in the human brain could have been attained. To address this question from a neurobiological perspective, we explored the association between the redundancy-to-synergy gradient and regional expression profiles of 20,674 genes from AIBS microarray data ^10,22^. Using partial least squares (PLS) regression, we show that the first two PLS components explained 31% of the variance in the regional synergy-redundancy values (Figure S12): significantly more than could be expected by chance (permutation test, p=0.007). For both components, gene expression weights were positively correlated with the redundancy-to-synergy regional gradient (PLS1: ρ = 0.37, p<0.001; PLS2: ρ = 0.39, p<0.001; Figure 4A and Figure S13). These correlations indicate that a number of genes are overexpressed in regions where synergy dominates over redundancy -- including significant overexpression of HAR-Brain genes, in line with the results presented above (PLS1: p=0.022; PLS2: p<0.001; Figure S14).

**Figure 4.**
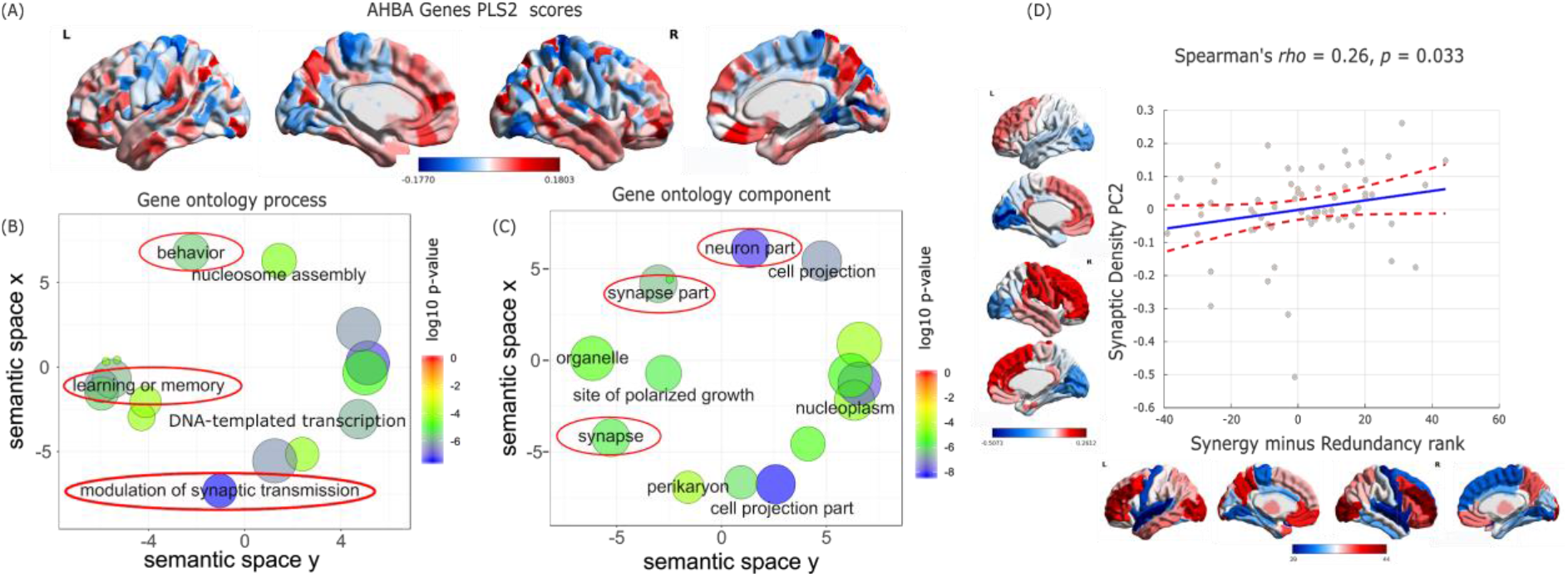
Neurobiological underpinnings of synergy in the human brain. (A) Second principal component of PLS (PLS2) relating the redundancy-to-synergy regional gradient to 20,647 genes from the Allen Institute for Brain Science, for the 308-ROI subdivision of the Desikan-Killiany cortical parcellation. (B) Dimensionality-reduced gene ontology terms pertaining to biological processes that are significantly enriched in PLS2 (red ovals highlight psychologically- or neurobiologically-relevant terms). (C) Dimensionality-reduced gene ontology terms pertaining to cellular components that are significantly enriched in PLS2 (red ovals highlight psychologically- or neurobiologically-relevant terms). Note that semantic space axes indicate the relative distance between terms in multi-dimensional space, but have no intrinsic meaning. Corresponding gene ontology terms for PLS1 are shown in Figure S15. (D) Significant correlation between regional redundancy-to-synergy gradient scores and an anterior-posterior principal component of synaptic density from [^11^C]UCB-J PET, for the DK-66 cortical parcellation. Corresponding results for the first principal component of [^11^C]UCB-J binding potential are shown in Figure S16.

We next sought to identify the role played by overexpressed genes related to brain synergy, for each PLS component. Analysis of gene ontology revealed that the transcriptional signature of PLS2 was significantly enriched in genes involved in learning or memory (in line with our meta-analytic results from NeuroSynth), as well as synapses, synapse components and synaptic transmission (all p<10^−4^ for significant enrichment).

Synapses are the key structures by which neurons exchange information; therefore they constitute a prime candidate for the neurobiological underpinning of synergistic interactions in the human brain, as suggested by our genetic analysis. To provide a more direct link between synaptic density and regional prevalence of synergy, we used positron emission tomography (PET) to estimate in vivo regional synaptic density based on the binding potential of the synapse-specific radioligand [^11^C]UCB-J ^23^. This radioligand has high affinity for the synaptic vesicle glycoprotein 2A (SV2A) ^24^, which is ubiquitously expressed in all synapses throughout the brain ^25^. Supporting the notion that regional brain synergy is related to underlying synaptic density, we found that an anterior-posterior principal component of synaptic density derived from [^11^C]UCB-J PET is significantly correlated with the regional gradient from redundancy to synergy (ρ = 0.26, p = 0.033; Figure 4D).

Therefore, genetic and molecular evidence converge to indicate synapses and synaptic transmission as key neurobiological underpinnings of synergy in the brain - in line with the notion that synergy quantifies information integration, and its role in supporting higher cognition.

Decomposing interactions between brain regions into synergistic and redundant components illuminates how the brain addresses the inherent trade-off between robustness and integration, providing powerful insights that are beyond traditional methods of studying brain interactions (e.g. FC). Having demonstrated the crucial role of synergistic interactions in human cognitive architecture via meta-analytic and graph-theoretical approaches, we proceeded to identify their neurobiological underpinnings by combining genetic, molecular and neuroanatomical evidence.

Taken together, our findings reveal that basic sensorimotor functions are supported by a modular backbone of redundant interactions (Fig 1D, 2B). As the brain’s input-output systems, reliable sensorimotor channels are vital for survival, warranting the additional robustness provided by redundant interactions — as indicated by our structural-functional analysis (Fig. 2D). In contrast, synergistic interactions are ideally poised to act as a global workspace, allowing the integration of complementary information from across the brain in the service of higher cognitive functions (Fig 1D): they bridge across different modules (Fig 1C), form a globally efficient network (Fig 2A), and their neuroanatomical organisation coincides with synapse-rich association cortex (Fig 4D and Supplementary Fig 3).

We further discovered that synergistic interactions were specifically enhanced in humans as a result of evolutionary pressures, with dedicated cortical real estate and dedicated genes, including those promoting synaptic transmission. This process resulted in a neural architecture that is capable of leveraging synergistic information to a greater extent than other primates. Our findings suggest that regions of the default mode and executive control (sub)networks may be able to support human higher cognition precisely thanks to their extensive involvement with synergistic processing.

Intriguingly, the high-synergy DMN is involved in self-related cognitive processes ^26,27^, and it is also especially disrupted by loss of consciousness, whether caused by anaesthesia or severe brain injury ^28^. Indeed, the global workspace theory of consciousness posits that integration of information within a global workspace is necessary for consciousness ^29^ - and a formal link has also been established between synergy and the measure of consciousness known as integrated information ^3,30^. Therefore, decomposition of information exchange into synergy and redundancy may also shed light on the emergence of consciousness in the human brain – providing a framework to discover the information-processing principles that govern how mental phenomena emerge from neurobiology.

## MATERIALS AND METHODS

### Synergy and Redundancy calculation

Shannon’s Mutual information (MI) quantifies the interdependence between two random variables *X* and *Y*. It is calculated as

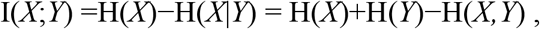

where H(*X*) stands for the Shannon entropy of a variable *X*. Above, the first equality states that the mutual information is equal to the reduction in entropy (i.e. uncertainty) about *X* after *Y* becomes accessible. Put simply, the mutual information quantifies the information that one variable provides about another ^31^.

Crucially, Williams and Beer (2010) ^2^ observed that the information that two source variables *X* and *Y* give about a third target variable *Z*, I(*X,Y* ; *Z*), should be decomposable in terms of different *types* of information: information provided by one source but not the other (unique information), or by both sources separately (redundant information), or jointly by their combination (synergistic information). Following this intuition, they developed the Partial Information Decomposition (PID ^2^) framework, which leads to the following fundamental decomposition:

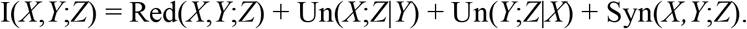

Above, *Un* corresponds to the unique information one source but the other doesn’t, *Red* is the redundancy between both sources, and *Syn* is their synergy: information that neither *X* nor *Y* alone can provide, but that can be obtained by considering *X* and *Y* together. It is worth noticing that the unique information is fully determined after synergistic and redundant comments have been accounted for; hence, we focus our analyses on the two latter components.

The simplest example of a purely synergistic system is one in which *X* and *Y* are independent fair coins, and *Z* is determined by the exclusive-OR function *Z* = XOR(*X,Y*): i.e., *Z*=0 whenever *X* and *Y* have the same value, and *Z*=1 otherwise. It can be shown that *X* and *Y* are both statistically independent of *Z*, which implies that neither of them provide - by themselves - information about *Z*. However, *X* and *Y* together fully determine *Z*: hence, the relationship between *Z* with *X* and *Y* is purely synergistic.

While PID provides a formal framework, it does not enforce how the corresponding parts ought to be calculated. While there is ongoing research on the advantages of different decompositions for discrete data, most decompositions converge into the same simple form for the case of continuous Gaussian variables ^32^. Known as *minimum mutual information PID* (MMI-PID), this decomposition quantifies redundancy in terms of the minimum mutual information of each individual source with the target; synergy, then, becomes identified with the additional information provided by the weaker source once the stronger source is known. Since linear-Gaussian models are sufficiently good descriptors of functional MRI timeseries (and more complex, non-linear models offer no advantage ^33^), here we adopt the MMI-PID decomposition, following previous applications of PID to neuroscientific data ^34^.

In a dynamical system such as the brain, one can calculate the amount of information flowing from the system’s past to its future, known as time-delayed mutual information (TDMI). Specifically, by denoting the past of variables as *X*_*t-τ*_ and *Y*_*t-τ*_ and treating them as sources, and their joint future state (*X*_*t*_, *Y*_*t*_), as target, one can apply the PID framework and decompose the information flowing from past to future as

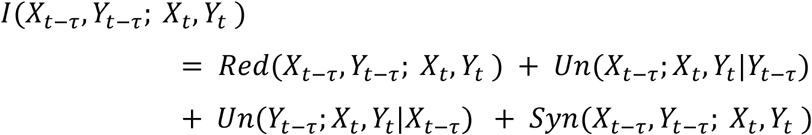

Recently, this equation has been refined to also distinguish between redundant, unique, and synergistic information shared with respect to the future variables *X*_*t*_, *Y*_*t*_. Importantly, this framework, known as Integrated Information Decomposition (PhiID) ^3^, has identified *Syn(X*_*t-τ*_, *Y*_*t-τ*_,*X*_*t*_,*Y*_*t*_) with the capacity of the system to exhibit emergent behaviour ^35^ [CITE emergence]. Furthermore, PhiID introduced a stronger notion of redundancy, in which information is shared by *X* and *Y* in both past and future. Accordingly, using the MMI-PhiID decomposition for Gaussian variables, we use

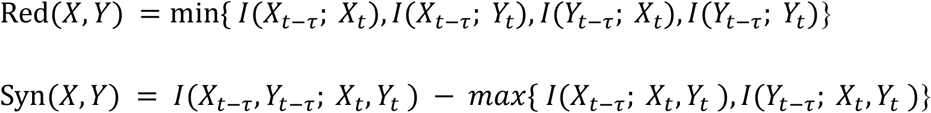

Here, we used the Gaussian solver implemented in the JIDT toolbox ^36^ to obtain TDMI, synergy and redundancy between each pair of brain regions, based on their HRF-deconvolved BOLD signal timeseries (Supplementary Methods).

### Gradient of redundancy-to-synergy relative importance

After building networks of synergistic and redundant interactions between each pair of regions of interest (ROIs), we determined the role of each ROI in terms of its relative engagement in synergistic or redundant interactions. We first calculated the nodal strength of each brain region as the sum of all its connections in the group-averaged matrix. Then, we ranked all 232 regions based on their nodal strength (with higher-strength regions having higher ranks). This procedure was done separately for networks of synergy and redundancy. Subtracting each region’s redundancy rank from its synergy rank yielded a gradient from negative (i.e. ranking higher in terms of redundancy than synergy) to positive (i.e. having a synergy rank higher than the corresponding redundancy rank); note that the sign is arbitrary.

It is important to note that the gradient is based on relative - rather than absolute - differences between regional synergy and redundancy. Consequently, a positive rank difference does not necessarily mean that the region’s synergy is greater than its redundancy; rather, it indicates that the balance between its synergy and redundancy relative to the rest of the brain is in favour of synergy - and *vice versa* for a negative gradient.

The same procedure was also repeated for network edges (instead of nodes), using their weights to rank them separately in terms of synergy and redundancy and then calculating their difference. This produced a single connectivity matrix where each edge’s weight represents its relative importance, being higher for synergy (positive edges) or redundancy (negative edges).

### NeuroSynth term-based meta-analysis of redundancy-to-synergy gradient

The regional redundancy-to-synergy gradient identified in terms of nodal rank differences was related to specific words using NeuroSynth, an online platform for large-scale, automated synthesis of fMRI data [https://neurosynth.org/]. For our analyses we employ 24 topic terms used by previous studies ^13,16^, which range from lower sensorimotor functions (such as eye movement, motion, visual and auditory perception) to higher cognitive functions (e.g. attention, working memory, social and numerical cognition).

A meta-analysis analogous to the one implemented by previous studies ^13,16^, was conducted to identify topic terms associated with the redundancy-to-synergy gradient. Twenty binary brain masks were obtained by splitting the values of the redundancy-to-synergy gradient into five-percentile increments. These brain masks served as input for the meta-analysis, based on the chosen 24 topic terms. For visualisation, terms were ordered according to the weighted mean of the resulting Z-statistics. Note that the term “visual semantics” was excluded from visualisation, because it failed to reach the significance threshold of Z > 3.1, leaving 23 terms (Figure 1). The analyses were carried out using modified code made freely available at [https://www.github.com/gpreti/GSP_StructuralDecouplingIndex].

### Measures of network integration and segregation

We quantified global integration in the networks of synergistic and redundant connections computing the networks *global efficiency*, a well-known measure that quantifies the ease of parallel information transfer in the network. More precisely, the global efficiency of a network corresponds to the average of the inverse of the shortest path length between each pair of nodes^37^

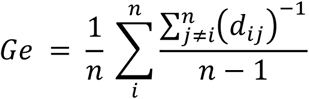

Following Cruzat et al (2018) ^38^, segregation of brain networks was quantified by means of network modularity. Put simply, the modularity function quantifies the extent to which a network can be partitioned such that the number of within-group edges is maximised and the density of between-group edges is minimised. We employed an implementation of Newman’s spectral modularity algorithm ^39^ available in the Brain Connectivity Toolbox (BCT; ^37,40^.

### Structural-Functional Similarity

Matrices of synergy and redundancy were thresholded proportionally using the same network density as the structural connectivity matrix of the same subject. This procedure was selected in order to ensure that the same number of edges would be present in both matrices, so that the two matrices can be compared. Then, the upper triangular portion of each connectivity matrix (structural and synergy/redundancy) was flattened into a vector, and the Spearman correlation coefficient between these two vectors was computed. We use this correlation as a measure of similarity between synergy or redundancy and structural connectivity.

### HAR-BRAIN genes

The maps of regional expression of human-accelerated genes for the DK-114 atlas were made available by Wei et al (2019), where the reader can find detailed information about how these data were generated. Briefly, genes located in a total of 2737 human accelerated regions (HARs) of the genome were taken as presented by comparative genome analysis representing genomic loci with accelerated divergence in humans ^41^. Out of 2143 HAR-associated genes identified from this procedure, 1711 were described in the Allen Human Brain Atlas (AHBA) microarray dataset (human.brain-map.org) ^21^ and were used in the analyses by Wei and colleagues, referred to as HAR genes.

HAR genes were subsequently subdivided into HAR-BRAIN and HAR-NonBRAIN genes. BRAIN genes were selected as the set of genes commonly expressed in human brain tissue using the Genotype-Tissue Expression (GTEx) database (data source: GTEx Analysis Release V6p; https://www.gtexportal.org/), which includes 56,238 gene expression profiles in 53 body sites collected from 7333 postmortem samples in 449 individuals. From these 56,238 genes, a total number of 2823 genes were identified as BRAIN genes showing significantly higher expressions in brain sites than non-brain sites (one-sided t-test and an FDR corrected q < 0.05 were used). HAR-BRAIN genes were identified as the 405 genes that overlapped between the 2823 BRAIN genes and the 1711 HAR genes, whereas the remaining HAR genes were labelled as HAR-NonBRAIN genes. Finally, the HAR gene expression data were mapped to the 114-region subdivision of the Desikan-Killiany atlas [DK-114] ^42,43^. Since only two of the six AHBA donors have data for the right hemisphere, Wei et al (2019) only considered HAR gene expression patterns for the left hemisphere.

### Cortical expansion

The maps of evolutionary cortical expansion were made available by Wei et al (2019), ^20^ who describe in detail how these data were generated. Briefly, Wei and colleagues analysed in-vivo MRI data from 29 adult chimpanzees, as well as 30 adult human subjects from the Human Connectome Project. Pial surface reconstructions of chimpanzee and human T1-weighted MRI scans (processed with FreeSurefer v5.3.0; https://surfer.nmr.mgh.harvard.edu/) were used for both vertex-to-vertex mapping across chimpanzee and humans and also for subsequent computation of region-wise expansion for cortical morphometry. A regional-level cortical surface area (Si) was computed by summing up face areas within each cortical region, for all regions of the DK-114 atlas ^42,43^. Normalized cortical area was obtained by dividing the regional area by the area of the whole cortex. Cortical expansion between every pair of chimpanzee and human subjects was calculated based on both the raw (“unadjusted”) and normalized (“adjusted”) cortical surface area by

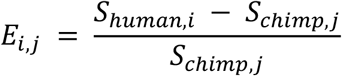

with E_i,j_ denoting the expansion from chimpanzee j to human i. A group-level region-wise cortical expansion map was calculated by taking averages over the 870 chimpanzee-to-human comparisons.

### AIBS gene expression analysis

Regional gene expression levels for 20,647 human genes were obtained from transcriptomic measurements in six post-mortem adult brains (age: 24-57 years), made available by the AIBS (human.brain-map.org) ^21^. We used code made freely available by Morgan et al (2019) ^10^ https://github.com/SarahMorgan/Morphometric_Similarity_SZ) to obtain a 308 x 20,647 regional transcription matrix, matching gene expression data to each cortical region of the DK-308 atlas ^10,22,44,45^ (Supplementary Methods). Each tissue sample was assigned to a cortical region using the AIBS MRI data for each donor, pooling samples between bilaterally homologous regions ^10,45^.

### Partial Least Squares

To explore the association between the redundancy-to-synergy regional gradient and all 20,647 genes measured in the AHBA microarrays, at each of 308 regions, we used partial least squares (PLS) as a dimensionality reduction technique ^10,22,44,46^. PLS finds components from the predictor variables (308 × 20,647 matrix of regional gene expression scores) that have maximum covariance with the response variables (308 × 1 matrix of regional redundancy-to-synergy gradient). The PLS components (i.e. linear combinations of the weighted gene expression scores) are ranked by covariance between predictor and response variables, so that the first few PLS components provide a low-dimensional representation of the covariance between the higher dimensional data matrices.

Goodness of fit of low-dimensional PLS components was tested non-parametrically by repeating the analysis 1000 times after shuffling the regional labels. The error on the PLS weights associated with each gene were tested by resampling with replacement of 308 ROIs (bootstrapping); the ratio of the weight of each gene to its bootstrap standard error was used to Z-score the genes and rank their contributions to each PLS component ^10,22,44^.

### Gene ontology and enrichment analysis

We used GOrilla for enrichment analysis of the first two PLS components ^22,47^ GOrilla identifies enriched gene ontology (GO) terms in ranked gene list, leveraging a large online database of gene annotations corresponding to ‘biological processes’ and ‘cellular components’ ^47^ We identified GO terms that were over-represented among the genes with the strongest positive weightings on each PLS component (i.e. those most strongly associated with dominance of synergy over redundancy). For our analyses on the online GOrilla platform (http://cbl-gorilla.cs.technion.ac.il) we unchecked the “Run GOrilla in fast mode” option and used the “P-value threshold 10-4” setting in order to best approximate FDR correction with α = 0.05 ^22^.

We then used the online tool REViGO (http://revigo.irb.hr) to summarize the list of significant GO terms and visualize the results of whole-genome enrichment analysis. First, REViGO employs measures of semantic similarity between terms ^48^ to identify representative clusters of genes. Then, REViGO plots significant GO terms in semantic space, where semantically similar GO terms are represented clustered near one another and labelled in a representative manner.

For our hypothesis-driven analysis, testing for enrichment of HAR-Brain genes, we also used non-parametric permutation testing. Specifically, we randomly drew 1000 samples of the same number of genes and estimated their PLS weighting, and compared the PLS weights of the HAR-Brain genes to this permutation distribution. This provided an estimate of the probability of HAR-Brain gene enrichment of each PLS component under the null hypothesis ^10,22^. We note that this permutation procedure does not take into account the correlation between HAR-Brain genes; more sophisticated null models for permutation testing that controlled for these or other characteristics of candidate genes will be important to develop for computational inference in future studies.

### Synaptic Density from Positron Emission Tomography

In-vivo estimates of regional synaptic density in the human brain were obtained from positron emission tomography (PET) with the radioligand [^11^C]UCB-J ((R)-1-((3-(methyl-^11^C)pyridin-4-yl)methyl)-4-(3,4,5-trifluorophenyl)pyr-rolidin-2-one) ^49^. This ligand quantifies synaptic density ^23^ based on its affinity for the presynaptic vesicle glycoprotein 2A (SV2A) ^24^ which is ubiquitously expressed in all brain synapses ^25^.

#### PET/MR imaging protocol

The research protocol was approved by an NHS Research Ethics Committee (REC: 18/EE/0059) and the Administration of Radioactive Substances Advisory Committee (ARSAC), and all participants provided written informed consent in accordance with the Declaration of Helsinki. Participant recruitment and exclusion criteria are described in detail in the original publication ^49^. Here, we included data from the healthy volunteers (N=15, 8 females; age: 68 ± 7 years).

The radioligand [^11^C]UCB-J was synthesised at the Radiopharmacy Unit, Wolfson Brain Imaging Centre, Cambridge University, using the methodology previously described ^24^. All participants underwent simultaneous 3T MRI and [^11^C]UCB-J PET on a GE SIGNA PET/MR (GE Healthcare, Waukesha, USA). Dynamic PET data acquisition was performed for 90 minutes starting immediately after [^11^C]UCB-J injection (median (range) injected activity: 408 (192-523) MBq, injected UCB-J mass ≤ 10 μg). Attenuation correction included the use of a multi-subject atlas method ^50^ and improvements to the MRI brain coil component ^51^. Each emission image series was aligned using SPM12 (www.fil.ion.ucl.ac.uk/spm/software/spm12/) then rigidly registered to a T1-weighted MRI acquired during PET data acquisition (TR = 3.6 msec, TE = 9.2 msec, 192 sagittal slices, in plane resolution 0.55 x 0.55 mm (subsequently interpolated to 1.0 x 1.0 mm); slice thickness 1.0 mm). Regional time-activity curves were extracted following the application of geometric transfer matrix partial volume correction ^51^ to each of the dynamic PET images. To quantify SV2A density (and therefore synaptic density), regional [^11^C]UCB-J non-displaceable binding potential (BP_ND_) was determined for a 66-ROI subdivision of the Desikan-Killiany cortical atlas (DK-66), using a basis function implementation of the simplified reference tissue model ^52^, with the reference tissue defined in the centrum semiovale ^53,54^.

#### Principal components of synaptic density

Principal Components Analysis (PCA) was subsequently employed to derive the principal components that explain most of the variance in regional [^11^C]UCB-J BP_ND_ across volunteers. Components were selected if their associated eigenvalue was greater than unity; two principal components satisfied this criterion, explaining 45% and 16% of the variance, respectively.

## Funding

This work was supported by grants from the National Institute for Health Research (NIHR, UK), Cambridge Biomedical Research Centre and NIHR Senior Investigator Awards [to DKM]; the British Oxygen Professorship of the Royal College of Anaesthetists [to DKM]; the Stephen Erskine Fellowship (Queens’ College, Cambridge), [to EAS]; and the Gates Cambridge Trust (to AIL). PAM and DB are funded by the Wellcome Trust (grant no. 210920/Z/18/Z). FR is funded by the Ad Astra Chandaria foundation. Computing infrastructure at the Wolfson Brain Imaging Centre (WBIC-HPHI) was funded by the MRC research infrastructure award (MR/M009041/1).

The PET study was funded by the Cambridge University Centre for Parkinson-Plus; the National Institute for Health Research Cambridge Biomedical Research Centre (146281); the Wellcome Trust (103838) and the Association of British Neurologists, Patrick Berthoud Charitable Trust (RG99368).

Data were provided [in part] by the Human Connectome Project, WU-Minn Consortium (Principal Investigators: David Van Essen and Kamil Ugurbil; 1U54MH091657) funded by the 16 NIH Institutes and Centers that support the NIH Blueprint for Neuroscience Research; and by the McDonnell Center for Systems Neuroscience at Washington University.

For the macaque data, primary support for the work by Newcastle University was provided by Wellcome Trust (WT091681MA, WT092606AIA), National Centre for 3Rs (Project grant NC/K000802/1; Pilot grant NC/K000608/1), and BBSRC (grant number BB/J009849/1).

We express our gratitude to the Primate neuroimaging Data-Exchange (PRIME-DE) initiative, to the organizers and managers of PRIME-DE and to all the institutions that contributed to the PRIME-DE dataset (http://fcon_1000.projects.nitrc.org/indi/indiPRIME.html), with special thanks to the Newcastle team. We are also grateful to Anoine Grigis, Jordy Tasserie and Bechir Jarraya for their help with the Pypreclin code, and Rodrigo Romero-Garcia for generating and sharing the 500mm2 subparcellation of the DK atlas, and the corresponding Von Economo cytoarchitectonics map. We are also grateful to Yongbin Wei and colleagues for generating and making available the data pertaining to HAR genes and cortical expansion. We are grateful to UCB Pharma for providing the precursor for the radioligand used in PET imaging.

## Author Contributions

AIL: conceived the study; analysed data; wrote first draft of the manuscript. PAM: conceived the study; contributed to data analysis; reviewed and edited the manuscript. FR: contributed to data analysis; reviewed and edited the manuscript. NH: acquired PET data; reviewed PET analysis; reviewed the manuscript. TDF: preprocessed PET data; reviewed the manuscript. JOB: conceived the PET project; reviewed PET analysis; reviewed the manuscript. JBR: conceived the PET project; reviewed PET analysis; reviewed the manuscript. DKM: reviewed the manuscript. DB: conceived the study; reviewed and edited the manuscript. EAS: conceived the study; reviewed and edited the manuscript.

### Competing Interests

JBR serves as an associate editor to Brain, and is a non-remunerated trustee of the Guarantors of Brain and the PSP Association (UK). He provides consultancy to Asceneuron, Biogen and UCB and has research grants from AZ-Medimmune, Janssen and Lilly as industry partners in the Dementias Platform UK. All other authors declare no conflicts of interest.

## Data Availability

The HCP DWI data in SRC format are available online (http://brain.labsolver.org/diffusion-mri-data/hcp-dmri-data). The HCP fMRI data are available online (https://www.humanconnectome.org/study/hcp-young-adult/data-releases).

Macaque MRI data are available from the PRIMatE Data Exchange (PRIME-DE) through the Neuroimaging Informatics Tools and Resources Clearinghouse (NITRC; http://fcon_1000.projects.nitrc.org/indi/indiPRIME.html).

The PET data that support the findings of this study are available from author NH, upon reasonable request for academic (non-commercial) purposes.

The macaque connectome is available online on Zenodo: https://zenodo.org/record/1471588#.X2JCjdZuJPY

Cortical gene expression patterns were taken from the transcriptomic data of the Allen Human Brain Atlas (AHBA, http://human.brain-map.org/static/download).

Region-wise maps of chimpanzee-to-human cortical expansion and HAR gene expression are available as Supplementary Materials from Wei et al (2019) ^20^.

The NMT anatomical volume and associated probabilistic tissue segmentation maps (GM, WM and CSF) are freely available online: https://afni.nimh.nih.gov/pub/dist/atlases/macaque/nmt and http://github.com/jms290/NMT.

## Code Availability

The Java Information Dynamics Toolbox is freely available online: (https://github.com/jlizier/jidt).

The CONN toolbox is freely available online (http://www.nitrc.org/projects/conn).

DSI Studio is freely available online (www.dsi-studio.labsolver.org).

The Brain Connectivity Toolbox code used for graph-theoretical analyses is freely available online (https://sites.google.com/site/bctnet/).

The code used for NeuroSynth meta-analysis is freely available online: (https://www.github.com/gpreti/GSP_StructuralDecouplingIndex).

The HRF deconvolution toolbox is freely available online: (https://www.nitrc.org/projects/rshrf).

The Pypreclin pipeline code is freely available at GitHub (https://github.com/neurospin/pypreclin).

The code for PLS analysis of gene expression profiles is freely available online: https://github.com/SarahMorgan/Morphometric_Similarity_SZ.

## Notes

### Competing Interest Statement

The authors have declared no competing interest.

