## Supplementary Information for "A synergistic core for human brain evolution and cognition"

### Supplementary Materials and Methods

#### Human data from Human Connectome Project

##### HCP: dataset description.

The dataset of functional and structural neuroimaging data used in this work came from the Human Connectome Project (HCP, http://www.humanconnectome.org/), Release Q3. Per HCP protocol, all subjects gave written informed consent to the HCP consortium. These data contained fMRI and diffusion weighted imaging (DWI) acquisitions from 100 unrelated subjects of the HCP 900 data release [1]. All HCP scanning protocols were approved by the local Institutional Review Board at Washington University in St. Louis.

##### HCP: Functional data acquisition

The following sequences were used: Structural MRI: 3D MPRAGE T1-weighted, TR= 2400 ms, TE = 2.14 ms, TI = 1000 ms, flip angle = 8°, FOV= 224 × 224, voxel size = 0.7 mm isotropic. Two sessions of 15 min resting-state fMRI: gradient-echo EPI, TR= 720 ms, TE= 33.1 ms, flip angle = 52°, FOV= 208 × 180, voxel size = 2 mm isotropic. Here, we only used functional data from the first scanning session, in LR direction. HCP-minimally preprocessed images were used for all acquisitions [2].

**Functional MRI preprocessing and denoising**

We used the minimally preprocessed fMRI data from the HCP, which includes bias field correction, functional realignment, motion correction, and spatial normalisation to Montreal Neurological Institute (MNI-152) standard space with 2mm isotropic resampling resolution [2]. We also removed the first 10 volumes, to allow magnetisation to reach steady state. Additional denoising steps were performed using the CONN toolbox (http://www.nitrc.org/projects/conn), version 17f[3]. To reduce noise due to cardiac and motion artifacts, we applied the anatomical CompCor method of denoising the functional data. The anatomical CompCor method (also implemented within the CONN toolbox) involves regressing out of the functional data the following confounding effects: the first five principal components attributable to each individual's white matter signal, and the first five components attributable to individual cerebrospinal fluid (CSF) signal; six subject-specific realignment parameters (three translations and three rotations) as well as their first- order temporal derivatives [4]. Linear detrending was also applied, and the subject-specific denoised BOLD signal timeseries were band-pass filtered to eliminate both low-frequency drift effects and high-frequency noise, thus retaining frequencies between 0.008 and 0.09 Hz.

**HRF deconvolution**

Following previous work on the use of information-theoretic measures in the context of functional MRI data, we used state-of-the-art techniques [5] to deconvolve the hemodynamic response function from our regional BOLD signal timeseries.

The only exceptions were the computation of traditional functional connectivity (Pearson correlation - see below), for which the un-deconvolved BOLD signal timeseries were instead used, in line with common practice; and the comparison with macaque fMRI: since no HRF deconvolution was performed for macaque data, they were compared with non-deconvolved human data.

**HCP: Diffusion Weighted Data**

We used DWI data from the 100 unrelated subjects of the HCP 900 subjects data release [1]. The diffusion weighted (DW) acquisition protocol is covered in detail elsewhere [2].

The diffusion MRI scan was conducted on a Siemens 3T Skyra scanner using a 2D spin-echo single-shot multiband EPI sequence with a multi-band factor of 3 and monopolar gradient pulse. The spatial resolution was 1.25 mm isotropic. TR = 5500 ms, TE = 89.50 ms. The b-values were 1000, 2000, and 3000 s/mm2. The total number of diffusion sampling directions was 90, 90, and 90 for each of the shells in addition to 6 b0 images. We used the version of the data made available in DSI Studio-compatible format at https://pitt.app.box.com/v/HCP1065.

#### DWI reconstruction and fiber tracking

The minimally-preprocessed DWI data [2] were corrected for eddy current and susceptibility artifact. DWI data were then reconstructed using q-space diffeomorphic reconstruction (QSDR), as implemented in DSI Studio (www.dsi-studio.labsolver.org) [6]. QSDR is a model-free method that calculates the orientational distribution of the density of diffusing water in a standard space, to conserve the diffusible spins and preserve the continuity of fiber geometry for fiber tracking. QSDR first reconstructs diffusion-weighted images in native space and computes the quantitative anisotropy (QA) in each voxel. These QA values are used to warp the brain to a template QA volume in Montreal Neurological Institute (MNI) space using the statistical parametric mapping (SPM) nonlinear registration algorithm. A diffusion sampling length ratio of 2.5 was used, and the output resolution was 1 mm.

A modified FACT algorithm[7] was then used to perform deterministic fiber tracking on the reconstructed data, with the following parameters. Angular cutoff of 55$^{\circ}$, step size of 1.0 mm, minimum length of 10 mm, maximum length of 400mm, spin density function smoothing of 0.0, and a QA threshold determined by DWI signal in the CSF. Each of the streamlines generated was automatically screened for its termination location. A whole-brain white matter mask was created by applying DSI Studio's default anisotropy threshold (0.6 Otsu's threshold) to the SDF's anisotropy values. The mask was used to eliminate streamlines with premature termination in the white matter region. Deterministic fiber tracking was performed until 1,000,000 streamlines were reconstructed for each individual.

#### Macaque Data from PRIME-DE Initiative

The non-human primate MRI data were made available as part of the Primate neuroimaging Data-Exchange (PRIME-DE) monkey MRI data sharing initiative, a recently introduced open resource for non-human primate imaging [8].

##### Macaque dataset description

We used fMRI data from rhesus macaques (Macaca mulatta) scanned at Newcastle University. This samples includes 14 exemplars (12 male, 2 female); Age distribution: 3.9-13.14 years; Weight distribution: 7.2-18 kg (full sample description available online: http://fcon_1000.projects.nitrc.org/indi/PRIME/files/newcastle.csv and http://fcon_1000.projects.nitrc.org/indi/PRIME/newcastle.html).

Out of the 14 total animals present in the Newcastle sample, 10 had awake resting-state fMRI data; of these 10, all except the first had two scanning sessions available: to maximise our statistical power, these repeated sessions were included in the analysis. Thus, the total was 19 distinct sessions across 10 individual macaques.

*Ethics approval:* All of the animal procedures performed were approved by the UK Home Office and comply with the Animal Scientific Procedures Act (1986) on the care and use of animals in research and with the European Directive on the protection of animals used in research (2010/63/EU). We support the Animal Research Reporting of In Vivo Experiments (ARRIVE) principles on reporting animal research. All persons involved in this project were Home Office certified and the work was strictly regulated by the U.K. Home Office. Local Animal Welfare Review Body (AWERB) approval was obtained. The 3Rs principles compliance and assessment was conducted by National Centre for 3Rs (NC3Rs). Animal in Sciences Committee (UK) approval was obtained as part of the Home Office Project License approval.

*Animal care and housing:* All animals were housed and cared for in a group-housed colony, and animals performed behavioural training on various tasks for auditory and visual neuroscience. No training took place prior to MRI scanning.

#### Macaque MRI acquisition

Animals were scanned in a vertical Bruker 4.7T primate dedicated scanner, with single channel or 4-8 channel parallel imaging coils used. No contrast agent was used. Optimization of the magnetic field prior to data acquisition was performed by means of 2nd order shim, Bruker and custom scanning sequence optimisation.

Animals were scanned upright, with MRI compatible head-post or non-invasive head immobilisation, and working on tasks or at rest (here, only resting-state scans were included). Eye tracking, video and audio monitoring were employed during scanning.

Resting-state scanning was performed for 21.6 minutes, with a TR of 2600ms, 17ms TE, Effective Echo Spacing of 0.63ms, voxels size 1.22 x 1.22 x 1.24. Phase Encoding Direction: Encoded in columns. Structural scans comprised a T1 structural, MDEFT sequence with the following parameters: TE: 6ms; TR: 750 ms; Inversion delay: 700ms; Number of slices: 22; In-plane field of view: 12.8 x 9.6cm2 on a grid of 256 x 192 voxels; Voxel resolution: 0.5 x 0.5 x 2mm; Number of segments: 8.

#### Macaque functional MRI preprocessing and denoising

The macaque MRI data were preprocessed using the recently developed pipeline for non-human primate MRI analysis, *Pypreclin*, which addresses several specificities of monkey research. The pipeline is described in detail in the associated publication [9]. Briefly, it includes the following steps: (i) Slice-timing correction. (ii) Correction for the motion-induced, time-dependent B0 inhomogeneities. (iii) Reorientation from acquisition position to template; here, we used the recently developed National Institute of Mental Health Macaque Template (NMT): a high-resolution template of the average macaque brain generated from in vivo MRI of 31 rhesus macaques (Macaca mulatta) [10]. (iv) Realignment to the middle volume using FSL MCFLIRT function. (v) Normalisation and masking using Joe's Image Program (JIP) -align routine (http://www.nmr.mgh .harvard.edu/~jbm/jip/, Joe Mandeville, Massachusetts General Hospital, Harvard University, MA, USA), which is specifically designed for preclinical studies: the normalization step aligns (affine) and warps (non-linear alignment using distortion field) the anatomical data into a generic template space. (vi) B1 field correction for low-frequency intensity non-uniformities present in the data. (vii) Coregistration of functional and anatomical images, using JIP -align to register the mean functional image (moving image) to the anatomical image (fixed image) by applying a rigid transformation. The anatomical brain mask was obtained by warping the template brain mask using the deformation field previously computed during the normalization step. Then, the functional images were aligned with the template space by composing the normalization and coregistration spatial transformations.

Denoising: The aCompCor denoising method implemented in the CONN toolbox was used to denoise the macaque functional MRI data, to ensure consistency with the human data analysis pipeline. White matter and CSF masks were obtained from the corresponding probabilistic tissue maps of the high-resolution NMT template (eroded by 1 voxel); their first five principal components were regressed out of the functional data, as well as linear trends and 6 motion parameters (3 translations and 3 rotations) and their first derivatives.

Following previous work on macaque functional MRI [11], data were bandpass-filtered in the range of 0.0025-0.05 Hz. When comparing directly between human and macaque data, results were also replicated using the same bandpass filter of 0.008-0.09Hz used for human data.

#### Brain Parcellations

Human brains were parcellated into 232 cortical and subcortical regions of interest (ROIs). The 200 cortical ROIs were obtained from the scale-200 version of the recent local-global functional parcellation of Schaefer et al (2018) [12]. Since this parcellation only includes cortical regions, it was augmented with 32 subcortical ROIs from a recent subcortical functional parcellation [13]. We refer to this 232-ROI parcellation as the augmented “Schaefer-232” parcellation.

The data pertaining to regional PET-derived synaptic density, HAR gene expression, human cortical expansion, and Von Economo cytoarchitectonics were each only available according to specific parcellations: the Desikan-Killiany anatomical atlas with 66 cortical regions (DK-66) for PET data; a 114-ROI subparcellation of the Desikan-Killiany atlas for HAR genes and cortical expansion (DK-114); and a different subparcellation of the Desikan-Killiany atlas with 308 equally-sized ROIs of 500 mm2 each (DK-308 [14]), for the Von Economo cytoarchitectonic classes.

For each of these parcellations, we report the corresponding connectivity matrices for redundancy and synergy, as well as the redundancy-to-synergy gradient and associated NeuroSynth meta-analysis. For the DK-308 parcellation, we also report a replication of our results pertaining to network integration, segregation and structural-functional similarity.

Macaque functional data were parcellated according to the whole-cortex 82-ROI “Regional Mapping” (RM) atlas of Kotter and Wanke [15], nonlinearly registered to the NMT template used for preprocessing.

#### BOLD timeseries extraction

To construct matrices of functional connectivity, the timecourses of denoised BOLD signals were averaged between all voxels belonging to a given atlas-derived ROI, using the CONN toolbox. The resulting region-specific timecourses of each subject were then extracted for further analysis in MATLAB version 2016a.

#### Traditional Functional connectivity

For each pair of brain regions *i* and *j*, their traditional functional connectivity $FC_{ij}$ was computed as the Pearson correlation between their denoised BOLD signal timeseries.

#### Structural connectome construction

To construct matrices of structural connectivity, the edge weights $a_{ij}$ of the structural connectivity matrix $A$ were defined as the number of streamlines connecting end-to-end each of the regions in the atlas, normalised to lie between zero and one. Note that deterministic tractography produces naturally sparse matrices, so that no thresholding is required.

#### Von Economo cytoarchitectonic classes

Whitaker and Vertes (2016) [16] assigned each regions in the DK-308 cortical parcellation to one of the cytoarchitectural types delineated by von Economo. This atlas subdivided the cortex into five types according to the laminar structure of the cortex and roughly corresponding to functional cortical specializations. Briefly, the primary motor cortex/precentral gyrus are regions with poor laminar differentiation; two types of regions generally considered to be association cortices are then distinguished, as well as secondary and primary sensory areas. Since the original classification of structural types does not discriminate between true six-layered isocortex and mesocortex or allocortex, which have markedly different cytoarchitectures and ontogenies [17], two additional subtypes were added: limbic cortex (which included the entorhinal, retrosplenial, presubicular and cingulate cortices, and thus primarily constitutes allocortex); and the insular cortex, which contains granular, agranular and dysgranular regions, and is therefore not readily assigned a single structural type.

Subsequently, synergy and redundancy were obtained for the DK-308 parcellation as described above, as well as the redundancy-to-synergy gradient based on rank differences. The regional values of this gradient were then averaged across all ROIs belonging to each of the seven cytoarchitectonic classes. For each cytoarchitectonic class, a positive overall score indicates that cortical regions belonging to that class have overall higher importance for synergy than for redundancy - and vice-versa for negative scores.

#### Canonical resting-state subnetworks

We used the canonical subdivision of the brain into 7 cortical subnetworks of Yeo [18]: default mode (DMN), somatomotor (SOM), visual (VIS), salience/ventral attention (SAL), dorsal attention (DAN), fronto-parietal executive control (FPN) and limbic (LIM). Schaefer et al (2018) [12] assigned each ROI in their cortical parcellation to one of these canonical subnetworks. The 32 subcortical regions were all assigned to an 8th subcortical subnetwork (SUB). For the

DK-308 parcellation, ROIs were each assigned to the subnetwork of greatest overlap (with random assignment in case of a tie). As this atlas is a sub-parcellation of the DK-66 cortical atlas, no subcortical subnetwork was present.

#### Alternative measures of network integration and structural-functional similarity

As an alternative quantification of network integration, we also employed the measure of global integrative capacity developed by Cruzat and colleagues [19]. This measure is calculated by taking the un-thresholded connectivity as input, and rescaling it to the range of 0 to 1 by dividing it by its largest element (note that both synergy and redundancy are guaranteed to be non-negative, and therefore we did not need to take the absolute value). Then, the matrix is progressively thresholded with an increasing threshold ρ in 1% increments. At each threshold value, the size of the largest connected component of the resulting network is evaluated (normalised by the total number of nodes to lie between 0 and 1). The final measure of global integration is the integral of the curve of largest connected component sizes over all thresholds:

$$I_{GC} = \int_{\rho=0.01}^{0.99} GC(A_\rho)$$

where $GC(A_\rho)$ indicates the size (number of nodes) of the largest connected component of the network whose adjacency matrix A has been thresholded at threshold ρ, and N is the total number of nodes in the network. This procedure was applied for both redundancy and synergy of each subject.

As an alternative method to quantify the similarity of synergistic and redundant connections to the underlying structural connectome, we computed the Hamming distance between the connectivity patterns of each ROI in the functional and anatomical connectivity matrices [20]. The Hamming distance between binary vectors *a* and *b* is computed as the number of symbol substitutions required to turn one vector into the other (normalised by their length). The matrices were therefore binarised by setting all supra-threshold entries to unity, and all others to zero. In the present application, the Hamming distance measures the proportion of connections that need to be changed before the two connectivity patterns become the same.

This analysis was performed for each ROI in our augmented Schaefer-232 atlas; by averaging over all ROI values, we obtained a value of structural-functional connectivity distance. Both correlation and Hamming distance analysis were performed separately for synergy and redundancy matrices.

#### Replication of structural-functional similarity results in macaque brains

Individual structural connectomes were not available for the macaques included in this study. Anatomical connectivity for the macaque brain was instead obtained from the fully-weighted, whole-cortex macaque connectome recently developed by Shen and colleagues [21]. This connectome was generated by combining information from two different axonal tract-tracing studies from the CoCoMac database [http://cocomac.g-node.org/main/index.php?] with diffusion-based tractography obtained from 9 adult macaques (Macaca mulatta and Macaca fascicularis). The resulting connectome provides a matrix of weighted and directed anatomical connectivity between each of the 82 cortical ROIs of the RM atlas [15]. Since synergy and redundancy are undirected measures of connectivity, the matrix of directed anatomical connections was made undirected by averaging the strength of connections in the two directions.

As for humans, networks of synergistic and redundant interactions were thresholded proportionally using the same density as the anatomical connectivity matrix, to ensure that the same number of edges would be present in the two matrices being compared. Then, Spearman correlation was used as the measure of structural-functional similarity.

#### Human-Macaque Comparison of Synergy and Redundancy

Separately for humans and macaques, each subject's matrices of synergistic and redundant interactions were each divided by the corresponding subject's matrix of TDMI. The global mean of the resulting matrices (across rows and columns) therefore represents the proportion of total information exchange across the brain that is provided by synergistic (v. redundant) interactions.

The human data were parcellated for this analysis according to the 83-ROI Lausanne parcellation (corresponding to the original Desikan-Killiany atlas, plus subcortical regions [22]), thereby ensuring a similar number of ROIs in human (83) and macaque brains (82).

Since macaque brains were not HRF-deconvolved, for this analysis we also used synergy and redundancy obtained from non-deconvolved human fMRI. As shown above, the use of HRF deconvolution had negligible effects on synergy and redundancy calculations, arguably thanks to the high temporal resolution of HCP data.

To ensure that the observed differences in the proportion of synergistic information could not be attributed to differences in bandpass-filter, we also repeated this analysis with macaque data filtered in the range 0.008-0.09Hz (i.e. same range as the HCP human data).

#### Dynamic mean field model to control for differences in TR between human and macaque data

##### Model construction

To determine whether the differences in the proportion of synergy between humans and macaques could be driven by the different TR (0.72s for HCP data, and 2.6s for macaque data), we simulated human functional MRI data with the same TR as the macaque data, using a dynamic neuronal mean-field model derived from the collective behavior of empirically validated integrate-and-fire (80% excitatory and 20% inhibitory) neurons [23]. The model combines macroscale information about neuroanatomy and structural connectivity (from DTI) with excitatory and inhibitory neuronal populations interconnected by AMPA, NMDA and GABA synapses, providing a neurobiologically plausible account of regional neuronal firing rate. We set all model parameters to be the same as those used by Deco et al (2018) [23], except for the global coupling parameter G, which we fit (see below).

The structural connectivity for the model was obtained by following the procedure of Wang et al (2019) [24], which derives a consensus structural connectivity matrix *A* from the individual SC matrices of the 100 HCP subjects included in the present study. Hence, for each individual, the structural connectivity was obtained from deterministic tractography (as described above) performed with the Lausanne-83 parcellation. Subsequently, for each pair of regions *i* and *j*, if more than half of subjects had non-zero connection *i* and *j*, $A_{ij}$ was set to the average across all subjects with non-zero connections between *i* and *j*. Otherwise, $A_{ij}$ was set to zero.

A Balloon-Windkessel hemodynamic model was then used to turn simulated regional neuronal activity into simulated regional BOLD signal. The Balloon-Windkessel model considers the BOLD signal as a static nonlinear function of the normalized total deoxyhemoglobin voxel content, normalized venous volume, resting net oxygen extraction fraction by the capillary bed, and resting blood volume fraction. The BOLD-signal estimation for each brain area is computed from the level of neuronal activity in that particular area. Finally, simulated regional BOLD signal was bandpass filtered in the same range as the empirical data (0.008-0.09Hz).

#### Model fitting

The model has only one free parameter *G*, which scales the global coupling strength. To find the value of *G* that generates the most realistic data, we first generated 100 simulations with a TR of 0.72s (i.e. the same as the empirical HCP data) for each value of *G* between 0.1 and 2.5, using increments of 0.1. Following Deco et al (2018) [23], we selected the value of G that minimised the Kolmogorov-Smirnov distance between the empirical and simulated functional connectivity dynamics (FCD), which has been shown to provide a better fit than simply using the global functional connectivity. The KS distance between empirical and simulated FCD was minimised for a value of *G*=1.6 (Figure S9).

#### Simulated human fMRI data

To determine whether the observed difference in proportion of synergy between human and macaque brains could be explained exclusively by the difference in TR, we then generated another set of 100 simulations, using the empirically determined best-fitting *G* parameter of 1.6, but now with a TR of 2.6s, i.e. the same as the macaque data. The simulated data were filtered between 0.008-0.09Hz and their mean normalised synergy across all pairs of regions (i.e. the proportion of total information exchange accounted for by synergy) was compared with the normalised synergy empirically observed in macaque brains (Figure S10).

#### AIBS gene expression sample collections and processing

The Allen Human Brain Atlas is a publicly available online resource of microarray-based gene expression profiles for an anatomically comprehensive set of brain regions [25], made available by the Allen Institute for Brain Science (human.brain-map.org). The dataset is based on post-mortem tissue from 6 donors with no known history of neuropathological or neuropsychiatric disease, who also passed a set of serology, toxicology and RNA quality screens. The donors were a 24-year-old African American male (H0351.2001), a 39-year-old African American male (H0351.2002), a 57-year old Caucasian male (H0351.1009), a 31-year old Caucasian male (H0351.1012), a 49-year old Hispanic female (H0351.1015) and a 55-year old Caucasian male (H0351.1016).

The following steps were performed (http://human.brain-map.org). Each brain was cut into slabs (0.5-1.0 cm thick) and frozen. Slabs were sectioned to allow expert neuroanatomic annotation, delineation and sampling. RNA was isolated and microarray data were generated for about 500 samples per hemisphere, representing all anatomical structures in approximate proportion to their volume. Each sample is then associated with expression levels for about 60,000 gene probes with 93% of known genes represented by at least 2 probes. Expression data were averaged across all samples from all donors in the matching anatomical structure, across both hemispheres. The data were also averaged across probes corresponding to the same gene, excluding probes that were not matched to gene symbols in the AHBA data.

#### Statistical Analysis

One-sample non-parametric t-tests with 10,000 permutations were used to determine whether the synergy-redundancy scores were significantly different from zero for each of the Yeo resting-state subnetworks and for each cytoarchitectonic class of Von Economo; FDR correction for multiple comparisons was adopted according to the Benjamini-Hochberg procedure [26].

The statistical significance of within-group differences in network properties was determined with non-parametric permutation t-tests (repeated-measures), with 10,000 permutations. Between-subjects non-parametric t-tests (also with 10,000 permutations) were instead used to test the statistical significance of human-macaque comparisons. All tests were two-sided, with an alpha value of 0.05. The effect sizes were estimated using Hedges's *g*.

To ensure robustness to possible outliers, Spearman's rank-based correlation coefficient was used to quantify the association between regional gradients of redundancy-to-synergy scores, and other regional measures (HAR-Brain gene expression, cortical expansion, PLS components).

### Supplementary Discussion

Functional connectivity has been especially fruitful in the context of fMRI data, which provide whole-brain coverage with fine spatial resolution. However, it should be borne in mind that the BOLD signal is only an indirect and relatively sluggish measure of neuronal activity. Here, these concerns were minimised by the use of HCP data, which have high (subsecond) temporal resolution and a large number of timepoints, and have become the gold standard of modern fMRI research. Additionally, we deconvolved the hemodynamic response function from our data prior to analysis - although this step did not appear to affect our results - arguably due to the high temporal resolution of HCP data).

We decided to avoid the use of global signal regression as a denoising step, whose use is still subject of controversy: while GSR can be beneficial to mitigate motion artifacts, it may also contain neuronal signal of interest. Therefore, we instead adopted the alternative aCompCor denoising method [4].

We also acknowledge that although we used several parcellations, with different origins (functional or anatomical) and spatial granularity, none of them included the cerebellum, which deserves further investigation.

With respect to the PET-derived measure of synaptic density, we acknowledge that this method is indirect as [$^{11}$C]UCB-J non-displaceable binding potential ($BP_{ND}$) is a measure of SV2A density rather than synaptic density. Nonetheless, it is the only available method to estimate synaptic density in humans in vivo. We also acknowledge the potential for off-target binding, but preclinical data indicate very high correlations between UCB-J binding and synaptophysin, which is a marker of pre-synaptic vesicular density [27]. Binding potentials for SV2A radioligands such as [$^{11}$C]UCB-J can be confounded by the use of concurrent medication that may bind to SV2A. We did not enrol any individuals taking levetiracetam or any member of this family of drugs that are SV2A-specific ligands [28].

Kinetic analysis of [$^{11}$C]UCB-J PET data using arterial blood sampling was not carried out in this study; instead we used reference tissue modelling to reduce the demand on our cohort which focussed on dementia patients. Furthermore, reference tissue modelling of [$^{11}$C]UCB-J with the centrum semiovale as the reference tissue has been verified against arterial input function compartmental modelling in healthy controls [29,30].

The PET data were obtained from middle-aged to elderly volunteers who were recruited as healthy controls for patients with dementia [31], which may represent a potential confound when relating [$^{11}$C]UCB-J $BP_{ND}$ to the gradient of redundancy-to-synergy obtained from HCP young adult volunteers. However, this concern is mitigated by evidence that synaptic density is stable throughout the human adult lifespan, with only a slight decline after age 74 [32].

Several differences exist between the human and macaque data we used, including scanner field strength (3T vs 4.7T) and TR (0.72s vs 2.6s). However, we find it reassuring that even despite these differences, our results pertaining to network and structural properties of synergistic and redundant interactions were qualitatively the same in macaques and humans. Moreover, for the direct comparison between species we took several steps to account for potential confounds, repeating our analyses with macaque data filtered in the same range as the human data, and with non-deconvolved human data, to ensure robustness. Furthermore, we used simulated human data with the same TR as the empirical macaque data to show that the difference in brain synergy cannot be attributed to mere differences in TR. We also note that the quantities compared were the proportion of total information accounted for by synergy and redundancy, rather than absolute values, thus ensuring that values were in the same theoretical range in both species.

### Supplementary Figures

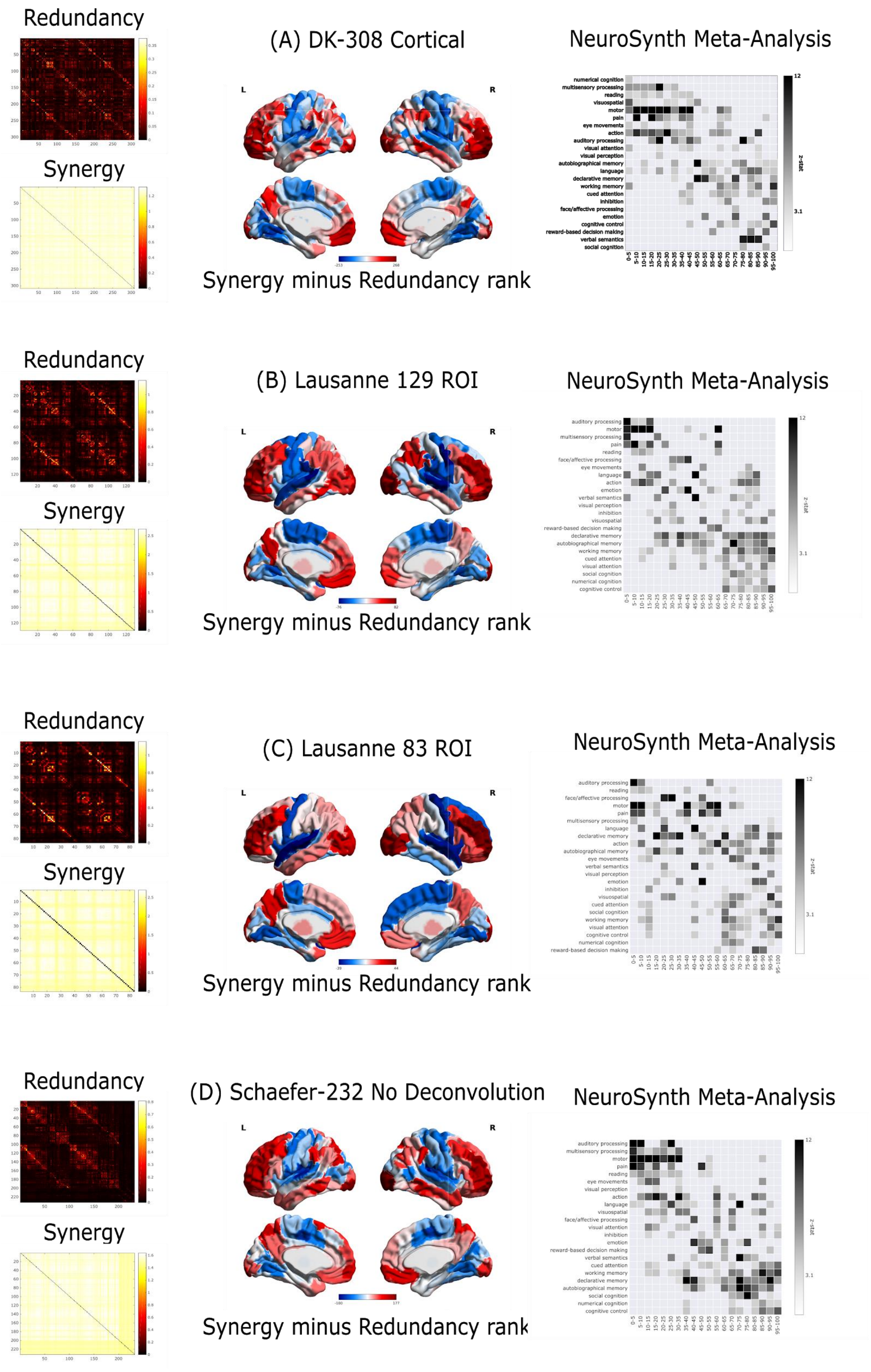
Redundancy
Synergy
(A) DK-308 Cortical
NeuroSynth Meta-Analysis
Synergy minus Redundancy rank
Redundancy
Synergy
(B) Lausanne 129 ROI
NeuroSynth Meta-Analysis
Synergy minus Redundancy rank
Redundancy
Synergy
(C) Lausanne 83 ROI
NeuroSynth Meta-Analysis
Synergy minus Redundancy rank
Redundancy
Synergy
(D) Schaefer-232 No Deconvolution
NeuroSynth Meta-Analysis
Synergy minus Redundancy rank

**Figure S1**. Replication of synergy-redundancy identification and NeuroSynth meta-analysis with alternative parcellations. Left: Group-average matrices of redundant and synergistic interactions; Middle: Redundancy-to-synergy gradient scores displayed on medial and lateral brain surfaces; Right: Results of the NeuroSynth term-based meta-analysis, relating the distribution of redundancy-to-synergy gradient across the brain to a gradient of cognitive domains, from lower-level sensorimotor processing to higher-level cognitive tasks (note that one term, "visual semantics", was excluded from visualisation because it failed to reach the threshold of Z > 3.1, leaving 23 terms). (A) DK-308 parcellation with equally-sized cortical areas (500 mm2), obtained as subdivisions of the Desikan-Killiany cortical atlas. (B) Lausanne-129 parcellation, comprising the DK-114 cortical ROIs, supplemented with 15 subcortical regions. (C) Lausanne-83 parcellation, comprising 68 cortical ROIs (Desikan-Killiany atlas), supplemented with 15 subcortical regions. (D) Schaefer-232 parcellation, without deconvolution of the hemodynamic response function (HRF) from the functional data.

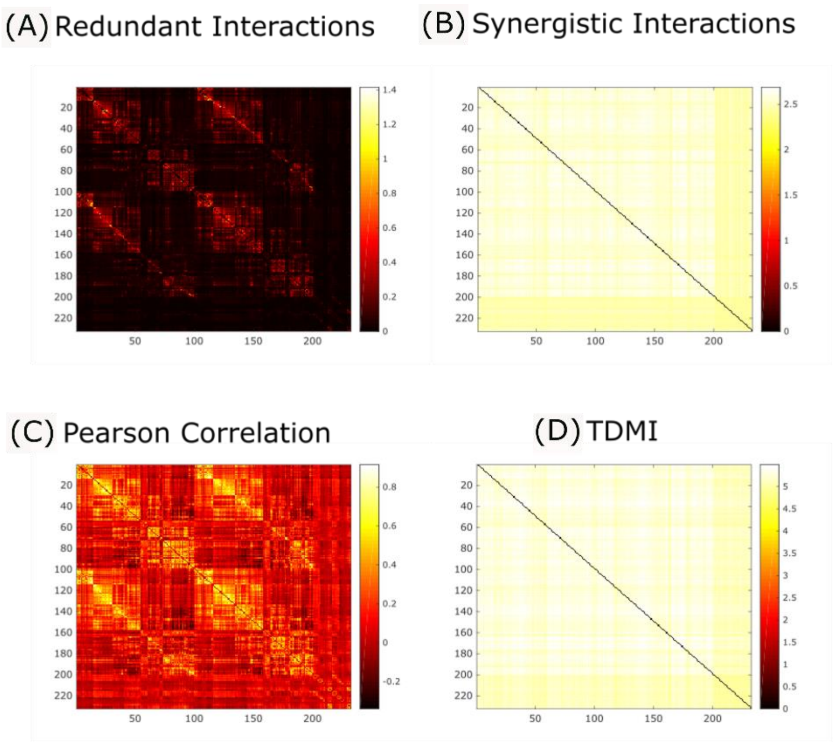


**Figure S2.** Group average of functional connectivity matrices obtained from (A) redundancy; (B) synergy; (C) Pearson correlation; and (D) time-delayed mutual information (TDMI).

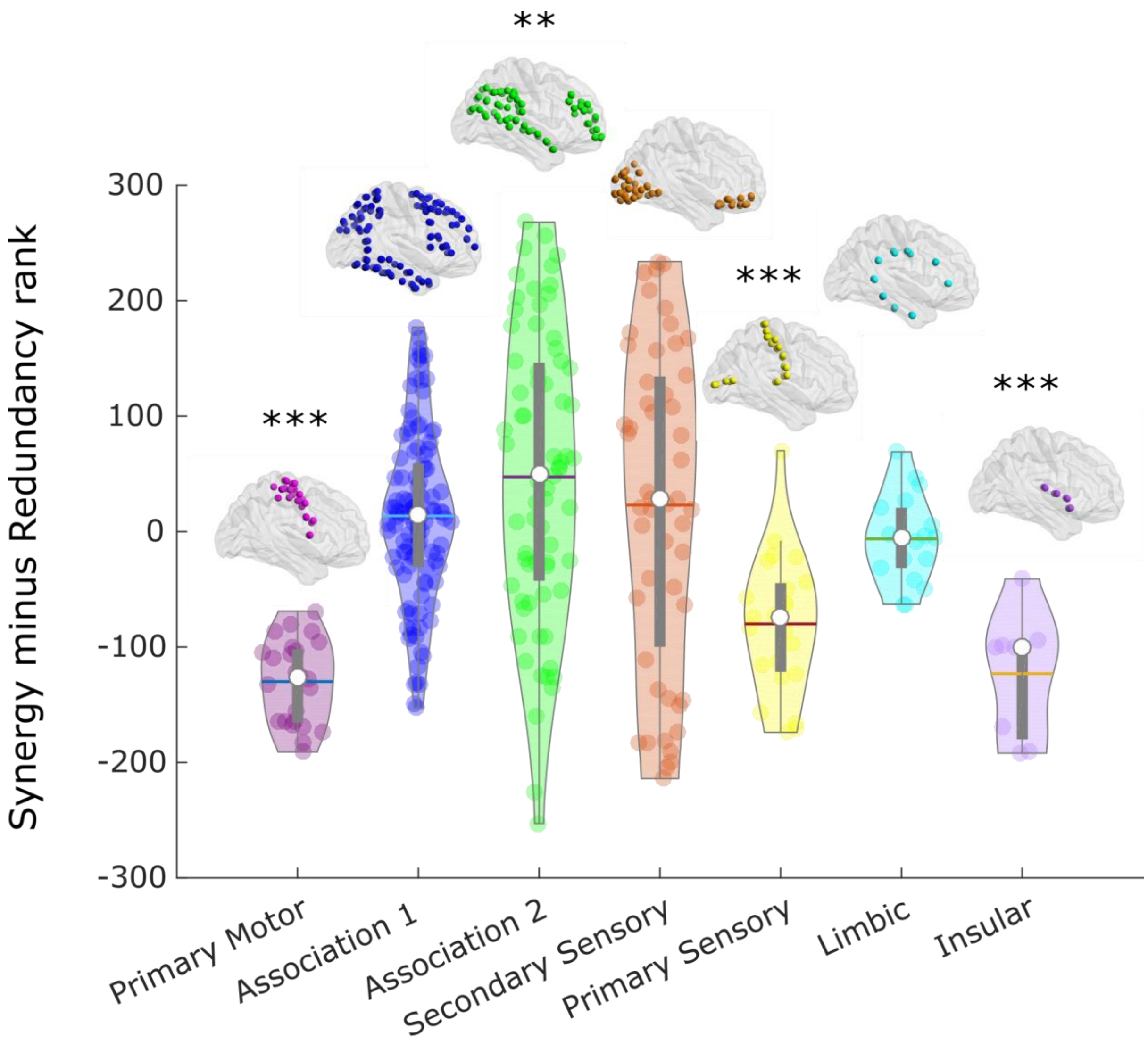


**Figure S3**. Regional redundancy-to-synergy gradient values for each of seven cytoarchitectonic classes (the five canonical classes identified by Von Economo, plus limbic and insular cortices), for 308 cortical ROIs of equal size (500 mm2), obtained from subdivisions of the Desikan-Killiany cortical parcellation. ** $p < 0.01$; *** $p < 0.001$, corrected for multiple comparisons using the False Discovery Rate.

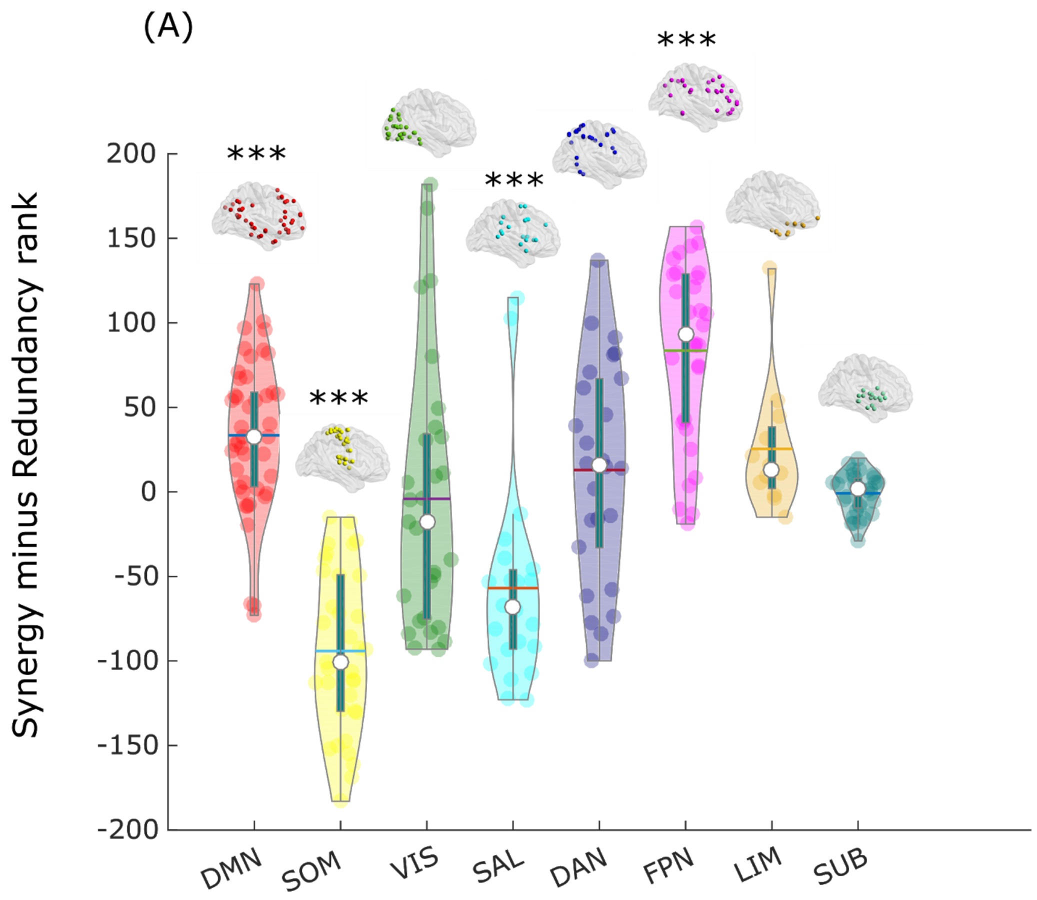
(A)
Synergy minus Redundancy rank
200
150
100
50
0
-50
-100
-150
-200
***
***
***
***
DMN
SOM
VIS
SAL
DAN
FPN
LIM
SUB

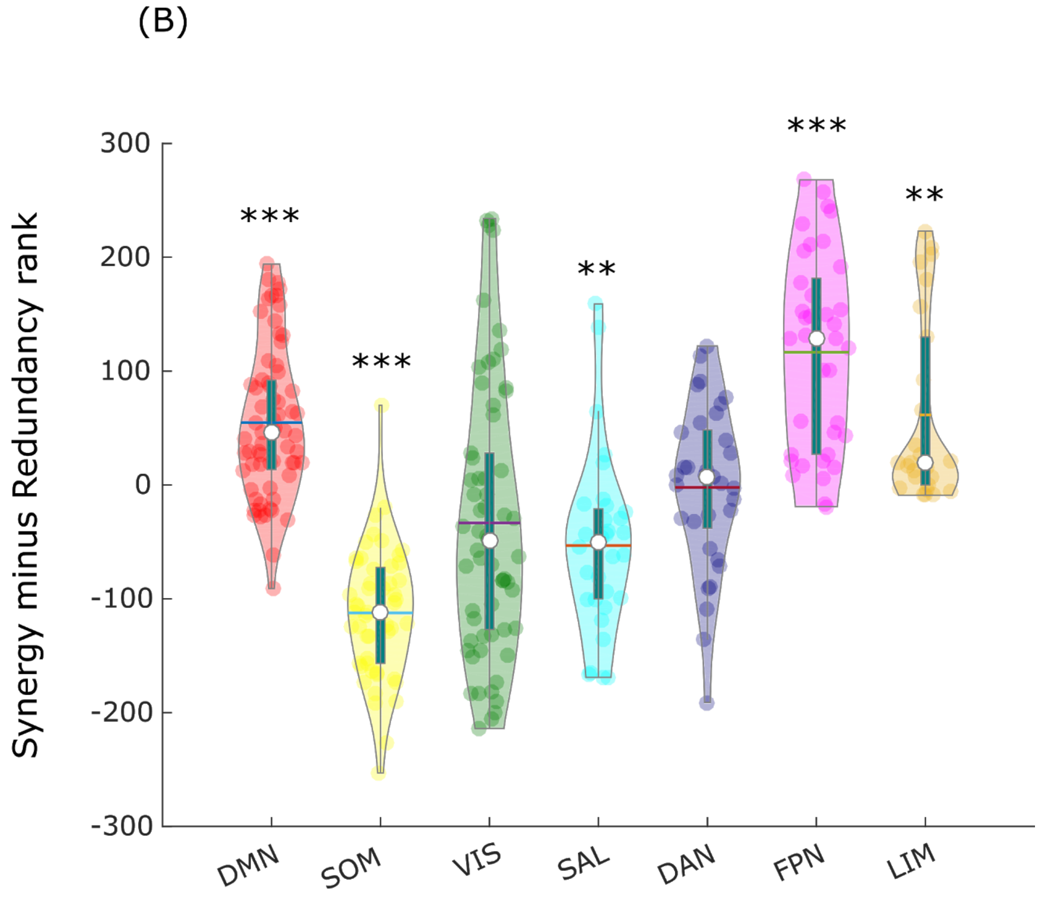
(B)
Synergy minus Redundancy rank
300
200
100
0
-100
-200
-300
***
***
**
***
**
DMN
SOM
VIS
SAL
DAN
FPN
LIM

464

**Figure S4**. Regional redundancy-to-synergy gradient values for each canonical resting-state network, as defined by Yeo et al (2011). (A) Schaefer-232 atlas, including 32 subcortical regions from the subcortical atlas of Tian et al (2020), which were assigned to an additional subcortical network. (B) 308 cortical ROIs of equal size (500 mm2), obtained from subdivisions of the Desikan-Killiany cortical parcellation. DMN, default mode network. SOM, somatomotor network. VIS, visual network. SAL, salience/ventral attention network; DAN, dorsal attention network. FPN, fronto-parietal executive control network. LIM, limbic network. SUB, subcortical network. ** $p < 0.01$; *** $p < 0.001$, corrected for multiple comparisons using the False Discovery Rate.

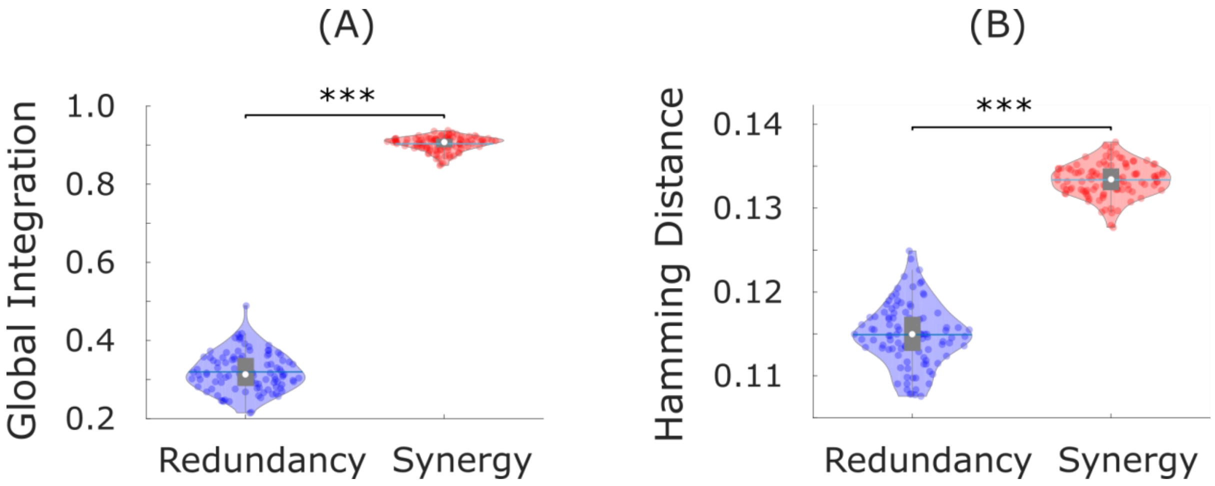


**Figure S5**. Significantly higher global integration (area under the curve of the size of the largest connected component across thresholds) and higher structural-functional dissimilarity (mean Hamming distance) for networks of synergy than redundancy. *** $p<0.001$.

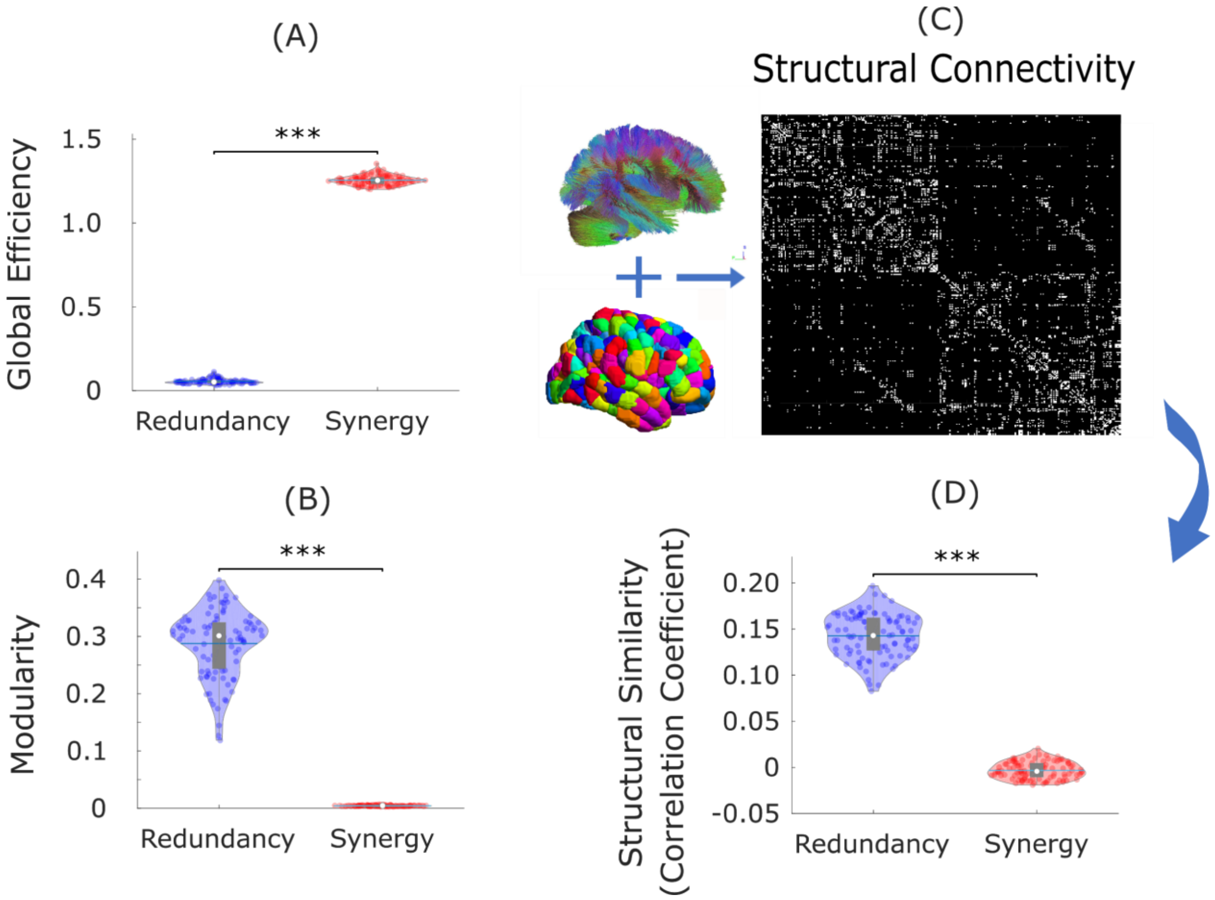


**Figure S6.** Replication of network results with 308-ROI cortical parcellation. (A) The network organisation of synergistic interactions exhibits significantly higher integrative capacity (global efficiency) than redundant interactions. (B) The network organisation of redundant interactions exhibits significantly higher segregation (modularity) than synergistic interactions. (C) Structural connectivity of each subject was estimated from diffusion MRI, measured as the number of white matter tracts between each pair of ROIs, and Spearman correlation coefficient was used to assess the similarity of redundancy and synergy matrices with structural connectivity, after thresholding to ensure equal numbers of connections. (D) Networks of redundant interactions are significantly more correlated with underlying structural connectivity than synergistic interactions. White circle: mean; blue line: median; *** $p < 0.001$.

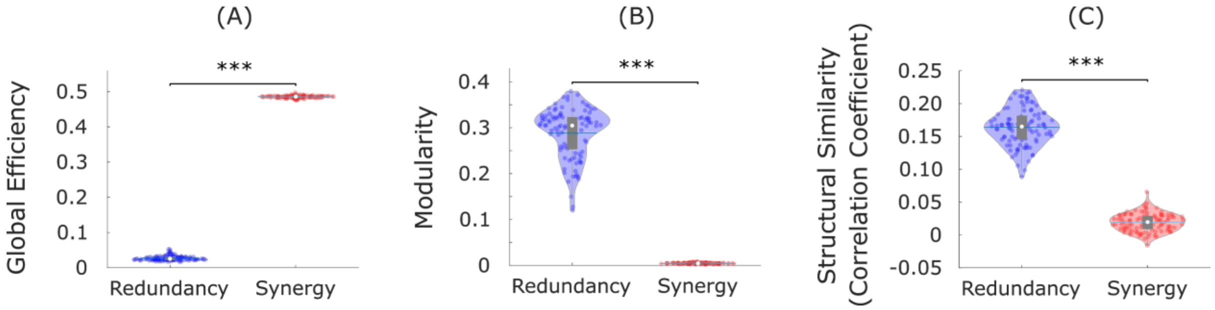


**Figure S7.** Replication of network results with synergy and redundancy normalised by TDMI. (A) The network organisation of synergistic interactions exhibits significantly higher integrative capacity (global efficiency) than redundant interactions. (B) The network organisation of redundant interactions exhibits significantly higher segregation (modularity) than synergistic interactions. (C) Networks of redundant interactions are significantly more correlated with underlying structural connectivity than synergistic interactions. White circle: mean; blue line: median; *** $p < 0.001$.

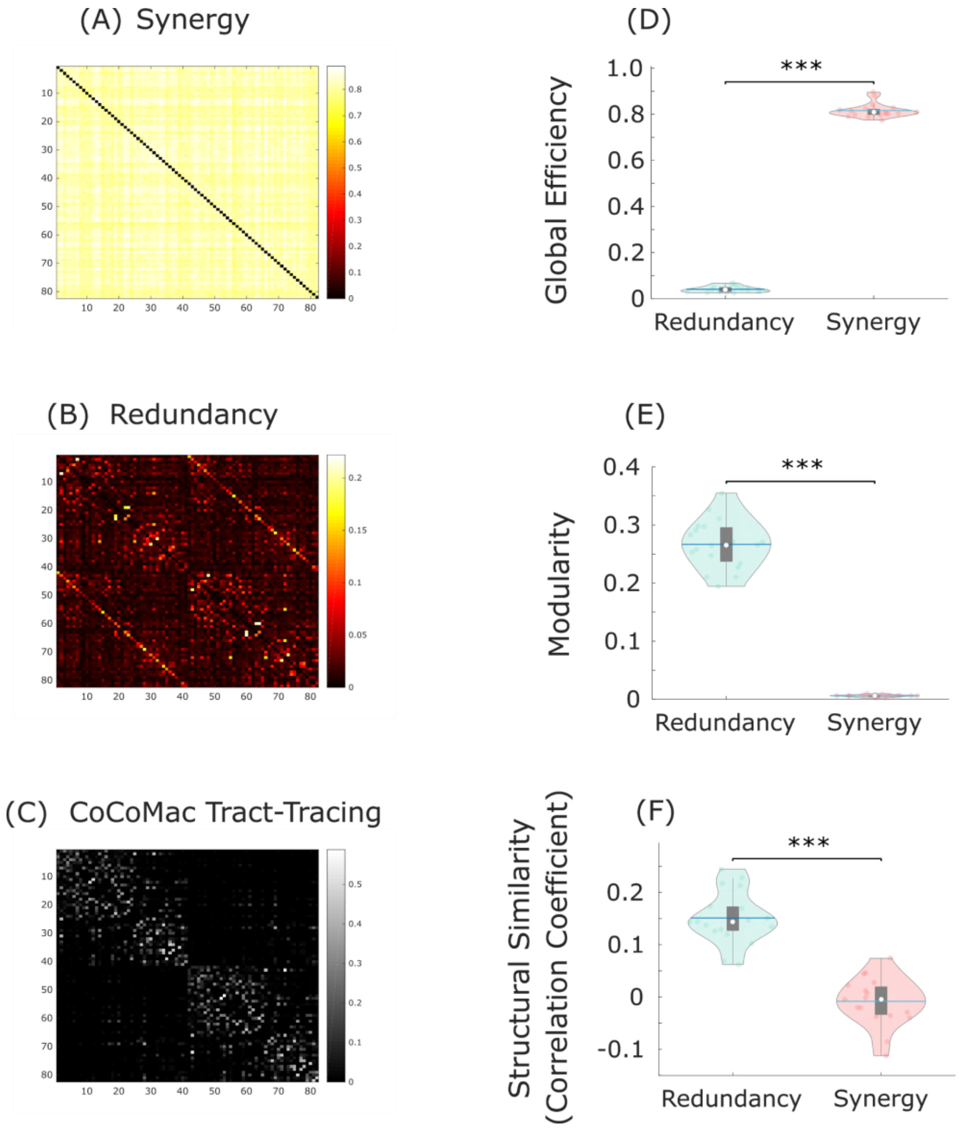

**Figure S8.** Characterisation of synergistic and redundant network profiles in macaque brains are similar to humans. (A) Synergistic interactions between regions of the macaque brain. (B) Redundant interactions between regions of the macaque brain. (C) Anatomical connectivity was estimated from axonal tracing (CIT), and Spearman correlation coefficient was used to assess the similarity of redundancy and synergy matrices with structural connectivity, after thresholding to ensure equal numbers of connections. (D) The network organisation of synergistic interactions exhibits significantly higher integrative capacity (global efficiency) than redundant interactions. (E) The network organisation of redundant interactions exhibits significantly higher segregation (modularity) than synergistic interactions. (F) Networks of redundant interactions are significantly more correlated with underlying anatomical connectivity than synergistic interactions. White circle: mean; blue line: median; *** $p < 0.001$.

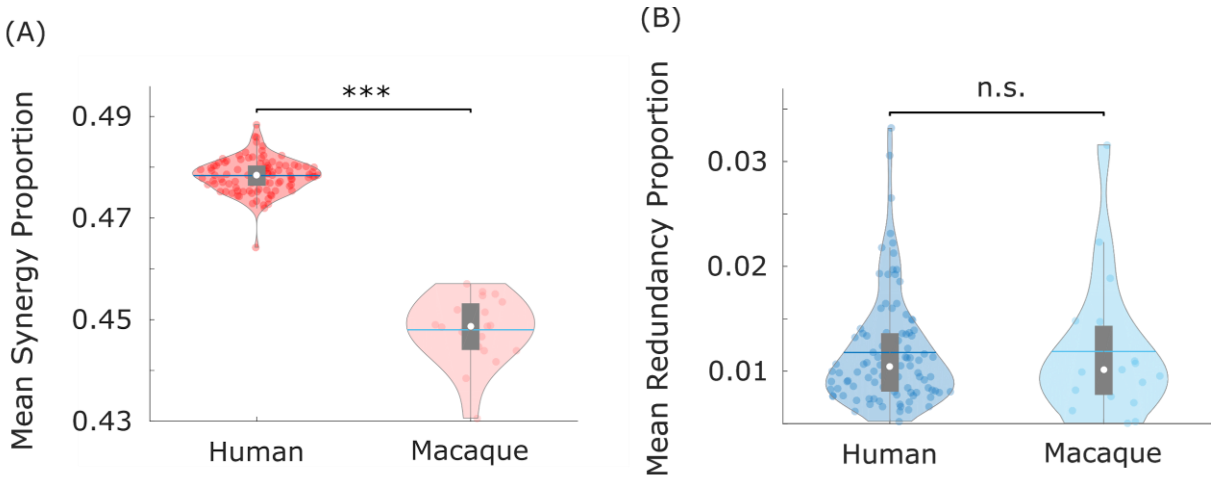


**Figure S9.** Replication of human-macaque comparison of synergy and redundancy proportion with same bandpass filtering. (A) The proportion of synergistic information exchange across the brain is significantly higher in humans (Homo sapiens) than monkeys (Macaca mulatta). (B) The proportion of redundant information exchange across the brain is equivalent in humans and macaques. To control for potential effects of bandpass filter, both human and macaque functional MRI data were bandpass filtered between 0.008-0.09Hz before calculation of synergy and redundancy values.

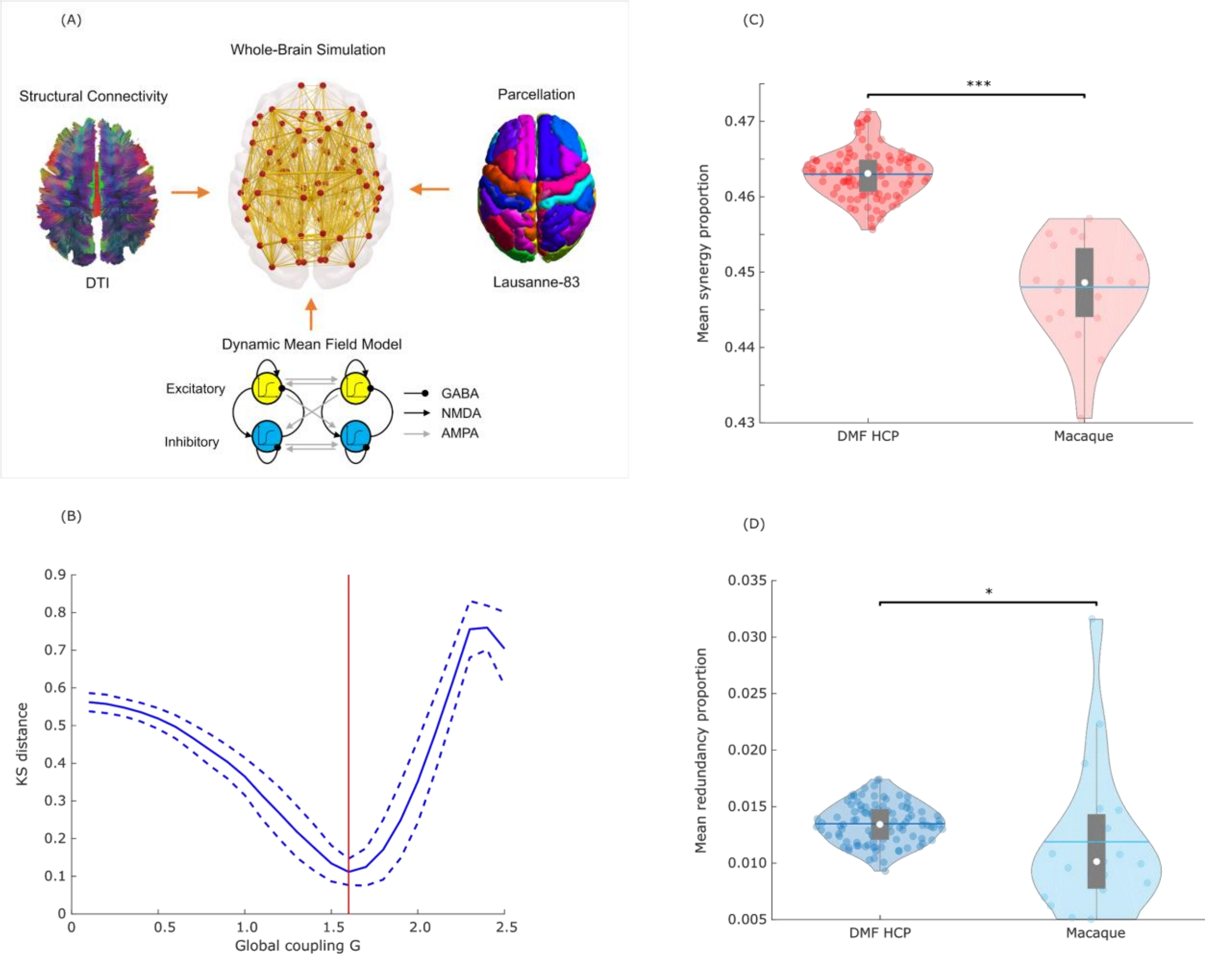


**Figure S10.** Simulation of human fMRI data with same TR as the macaque data shows that human-macaque differences in synergy cannot be attributed solely to TR differences between datasets. (A) The dynamic mean field (DMF) model used to simulate human fMRI data combines macroscale information about neuroanatomy and structural connectivity (from DTI) with excitatory and inhibitory neuronal populations interconnected by AMPA, NMDA and GABA synapses, providing a neurobiologically plausible account of regional neuronal firing rate, which is turned into simulated BOLD signal by means of the Balloon-Windkessel hemodynamic model. (B) Using a TR of 0.72s (the same as the empirical HCP data), the model is fitted to the empirical HCP data by finding the value of the global coupling parameter G that minimises the Kolmogorov-Smirnov distance between the distributions of empirical and simulated functional connectivity dynamics (FCD). The KS distance is minimised for G=1.6, which is the value of G used for subsequent simulations with TR=2.6s (the same TR as the macaque data). (C) The proportion of synergistic information exchange across the brain is significantly higher in simulated human data than in empirical macaque data with the same TR=2.6s ($p<0.001$). (D) The proportion of redundant information exchange across the brain is also significantly higher in simulated human data than empirical macaque data ($p=0.036$).

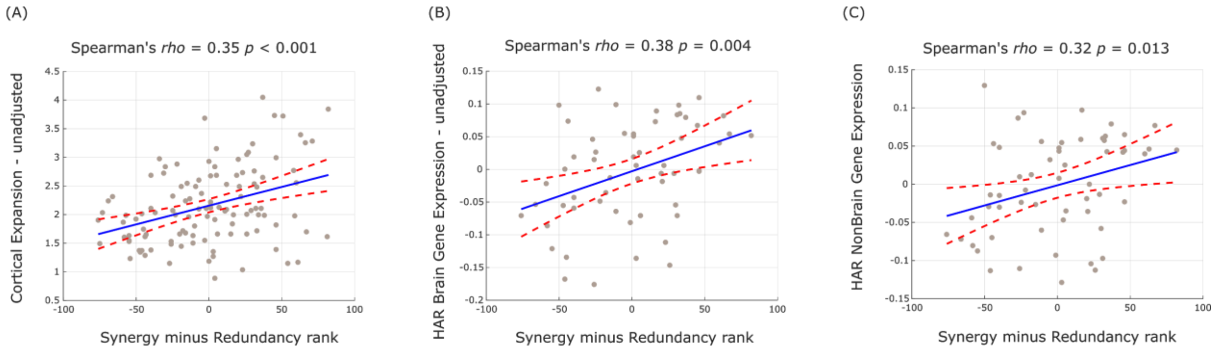


**Figure S11.** Synergy-redundancy gradient correlates with unadjusted cortical expansion and gene expression. (A) Significant correlation between regional redundancy-to-synergy gradient scores and unadjusted regional cortical expansion from chimpanzee (Pan troglodytes) to human (both on DK-114 cortical atlas, both hemispheres). (B) Significant correlation between regional redundancy-to-synergy gradient scores and unadjusted regional expression of brain-related human-accelerated (HAR) genes (both on left hemisphere of DK-114 atlas). (C) Significant correlation between regional redundancy-to-synergy gradient scores and unadjusted regional expression of non-brain-related human-accelerated (HAR) genes (both on left hemisphere of DK-114 atlas).

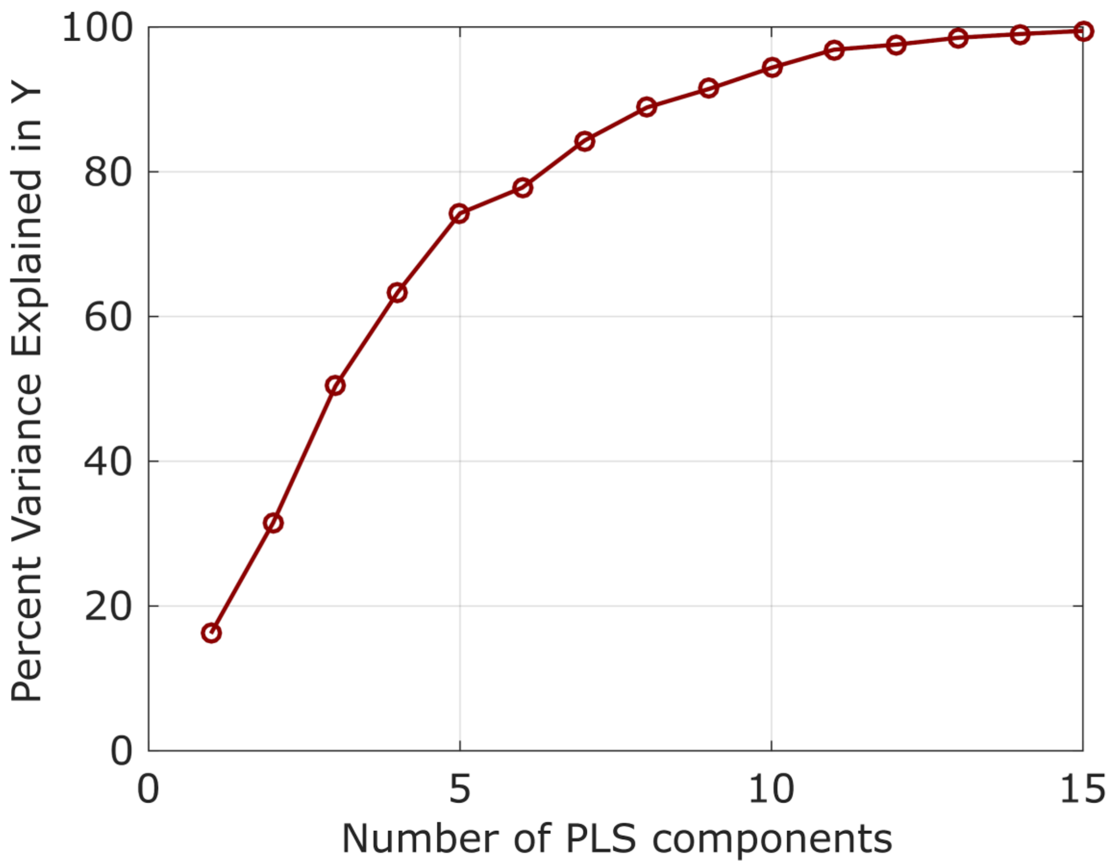


**Figure S12.** Percentage of variance explained in the pattern of regional redundancy-to-synergy scores, as a function of number of PLS components included.

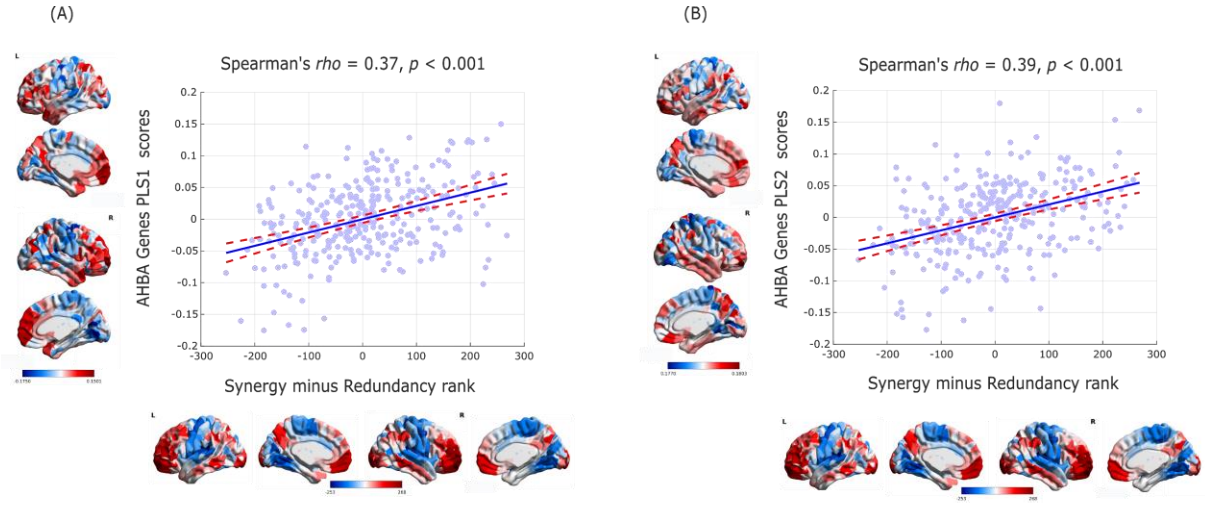


**Figure S13.** Correlations between PLS components of 20,647 genes from the Allen Institute for Brain Science and redundancy-to-synergy regional patterns, for the 308-ROI subdivision of the Desikan-Killiany cortical parcellation. (A) First principal component of PLS (PLS1). (B) Second principal component of PLS (PLS2).

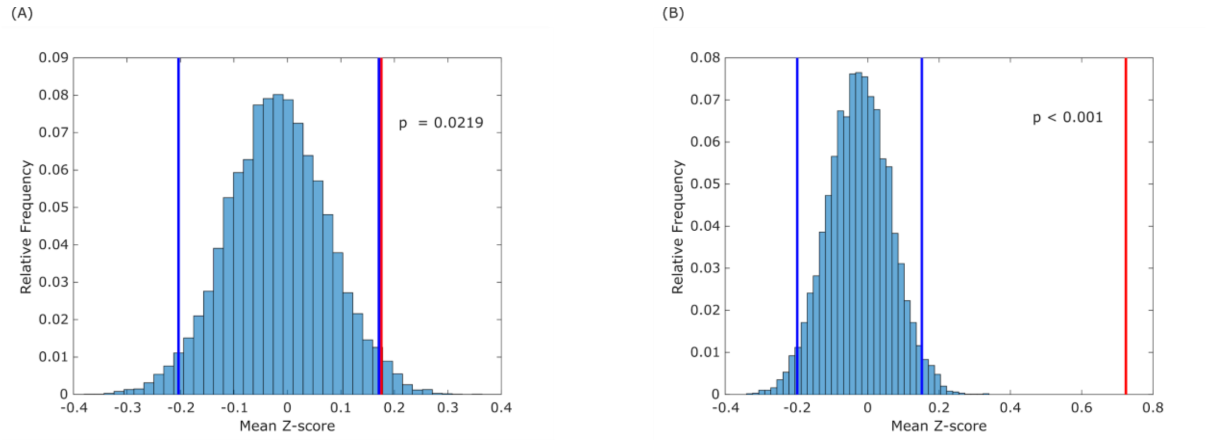


**Figure S14.** Significant enrichment of HAR-Brain genes in the first two components of PLS. (A) PLS1. (B) PLS2. Histograms indicate the relative frequency (over 1,000 bootstraps) of the mean Z-score of a random sample of genes of equal size as the HAR-Brain genes. Blue vertical lines: top and bottom 2.5% of the distribution of bootstrapped Z-scores. Red vertical line: empirical mean Z-score of HAR-Brain genes for the corresponding PLS component.

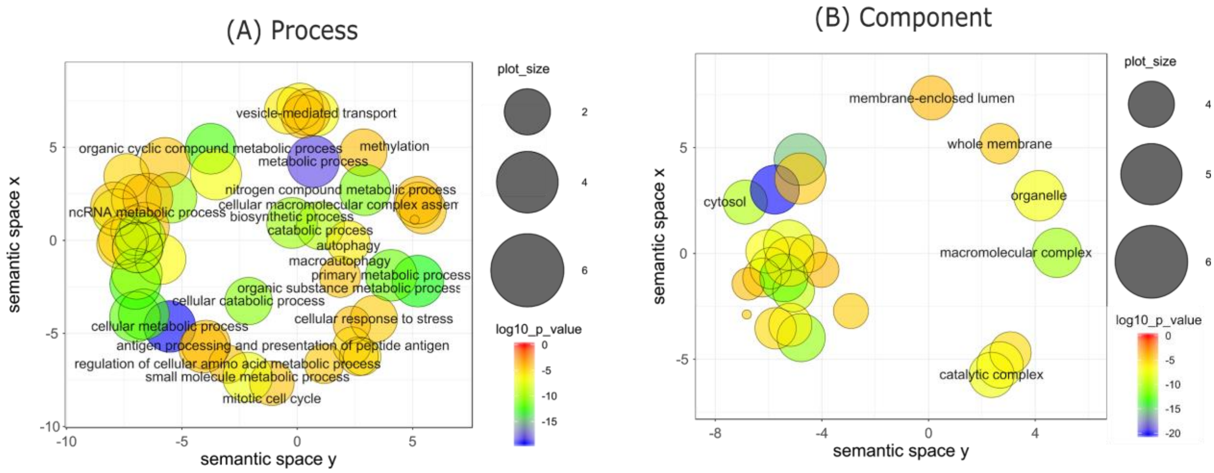


**Figure S15.** Gene ontology for PLS1. (A) Dimensionality-reduced gene ontology terms pertaining to biological processes that are significantly enriched in PLS1. (B) Dimensionality-reduced gene ontology terms pertaining to cellular components that are significantly enriched in PLS1. Note that semantic space axes indicate the relative distance between terms in multi-dimensional space, but have no intrinsic meaning.

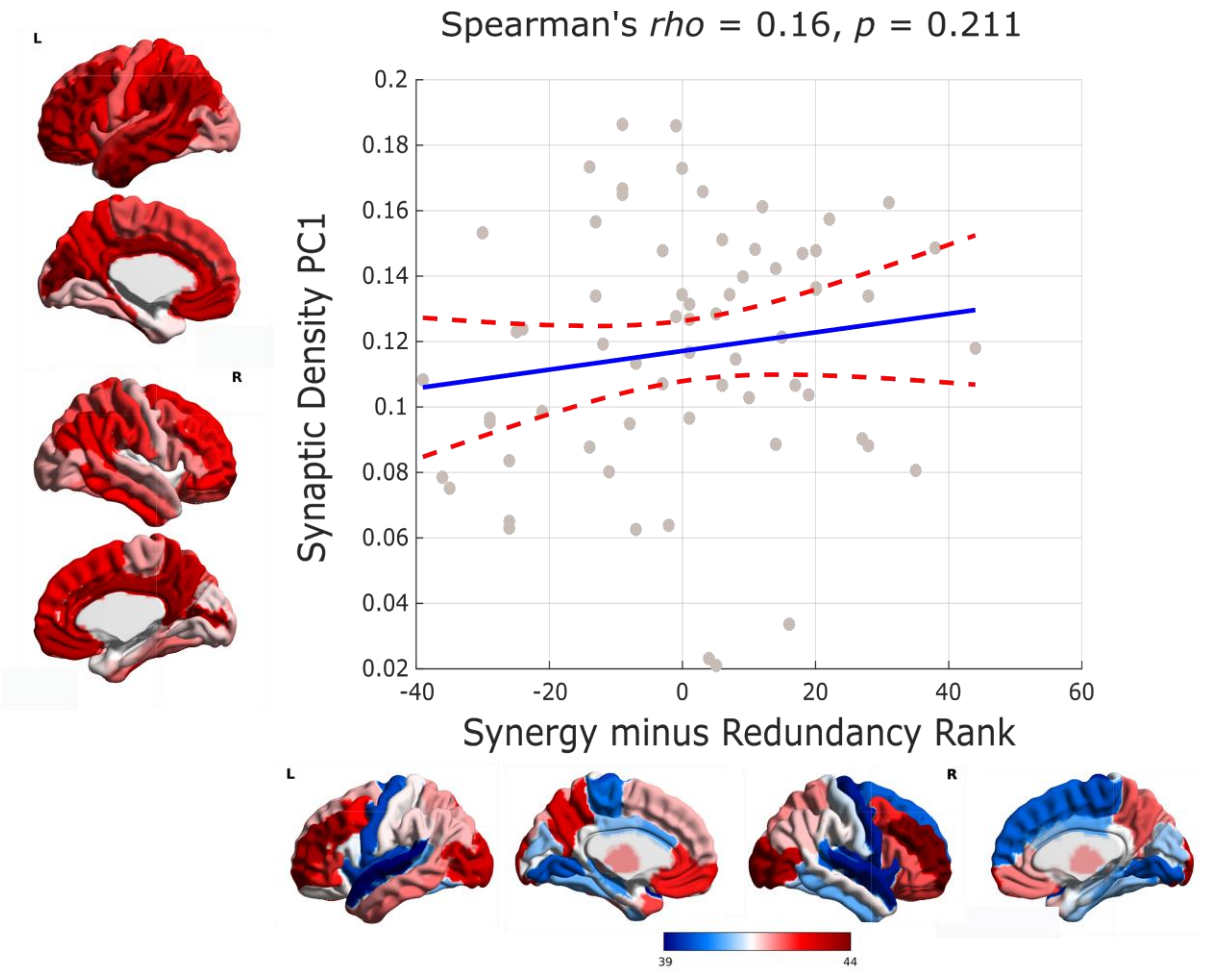

**Figure S16.** Non-significant correlation between regional redundancy-to-synergy gradient scores and first principal component of synaptic density from [[11]C]UCB-J binding potential. (Both on DK-66 atlas).

### Supplementary Tables

**Supplementary Table 1.** Synergy and redundancy network results for the augmented Schaefer-232 parcellation.

| Measure | Red Mean | Syn Mean | Red SD | Syn SD | t-score | df | Effect Size | p-value | Sig |
|---|---|---|---|---|---|---|---|---|---|
| Struct-Func Correlation | 0.16 | 0.02 | 0.03 | 0.01 | 39.85 | 99 | 6.29 | p<0.001 | *** |
| Modularity | 0.29 | 0.00 | 0.06 | 0.00 | 51.74 | 99 | 7.25 | p<0.001 | *** |
| Global Efficiency | 0.14 | 2.54 | 0.04 | 0.06 | -330.04 | 99 | -46.67 | p<0.001 | *** |
| Struct-Func Hamming Distance | 0.11 | 0.13 | 0.00 | 0.00 | -39.26 | 99 | -6.19 | p<0.001 | *** |
| Global Integration | 0.32 | 0.90 | 0.05 | 0.02 | -117.76 | 99 | -15.64 | p<0.001 | *** |

**Supplementary Table 2.** Synergy and redundancy network results for the 308-ROI 500mm2 cortical parcellation.

| Measure | Red Mean | Syn Mean | Red SD | Syn SD | t-score | df | Effect Size | p-value | Sig |
|---|---|---|---|---|---|---|---|---|---|
| Struct-Func Correlation | 0.14 | 0.00 | 0.02 | 0.01 | 54.14 | 99 | 8.18 | p<0.001 | *** |
| Modularity | 0.29 | 0.00 | 0.06 | 0.00 | 49.27 | 99 | 6.92 | p<0.001 | *** |
| Global Efficiency | 0.06 | 1.25 | 0.01 | 0.03 | -372.51 | 99 | -52.58 | p<0.001 | *** |

**Supplementary Table 3.** Synergy and redundancy network results for normalised synergy and redundancy (proportion of total information).

| Measure | Red Mean | Syn Mean | Red SD | Syn SD | t-score | df | Effect Size | p-value | Sig |
|---|---|---|---|---|---|---|---|---|---|
| Struct-Func Correlation | 0.164 | 0.019 | 0.028 | 0.013 | 49.635 | 99 | 6.574 | p<0.001 | *** |
| Modularity | 0.288 | 0.005 | 0.055 | 0.001 | 51.912 | 99 | 7.257 | p<0.001 | *** |
| Global Efficiency | 0.027 | 0.485 | 0.007 | 0.004 | -750.985 | 99 | -79.905 | p<0.001 | *** |

**Supplementary Table 4.** Synergy and redundancy network results for macaques.

| Measure | Red Mean | Syn Mean | Red SD | Syn SD | t-score | df | Effect Size | p-value | Sig |
|---|---|---|---|---|---|---|---|---|---|
| Struct-Func Correlation | 0.151 | -0.008 | 0.049 | 0.045 | 8.648 | 18 | 3.302 | p<0.001 | *** |
| Modularity | 0.267 | 0.006 | 0.041 | 0.002 | 27.419 | 18 | 8.638 | p<0.001 | *** |
| Global Efficiency | 0.041 | 0.816 | 0.013 | 0.030 | -136.378 | 18 | -32.906 | p<0.001 | *** |
